# Comparison of Activities of Transcription Factor NF-κB from Two Jellyfish Models

**DOI:** 10.1101/2025.01.21.634164

**Authors:** Leah M. Williams, Wei Wang, Alexandra V. Grigoryeva, Alejandra Navarro-Rosado, Jada A. Peart, Angela Calderon, Catherine L. Gill, Susan Black, Kristina M. Alsante, Aidan T. Lackstrom, Molecular Biology Laboratory, Brandon Weissbourd, Mengrui Wang, Christopher J. DiRusso, Lianne B. Cohen, Zeba Wunderlich, Brian P. Grone, Thomas D. Gilmore

## Abstract

Herein, transcription factor Nuclear Factor-kappaB (NF-κB) is characterized as the downstream effector of putative innate immune signaling pathways from two model jellyfish, *Aurelia aurita* (*Aa*) and *Clytia hemisphaerica* (*Ch*). Both jellyfish NF-κB proteins consist primarily of the N-terminal DNA-binding/dimerization domain, and they lack the C-terminal ankyrin repeat inhibitory domain found in vertebrate NF-κB proteins. Both jellyfish NF-κB proteins bind to a consensus mammalian NF-κB binding site, and their AlphaFold3-predicted structures on DNA are similar to that of mouse NF-κB p50. Neither *Aa*- nor *Ch*-NF-κB activated transcription of an NF-κB-site reporter gene in human cells; however, *Aa*-NF-κB did activate transcription from a GAL4-site reporter gene in yeast, whereas *Ch*-NF-κB did not. *Aa*- and *Ch*-NF-κB were both constitutively located in the nucleus when expressed in vertebrate cells. Homologs of IκB-like proteins from both species interacted with their corresponding NF-κB proteins in co-immunoprecipitation assays in HEK 293T cells. These IκB-like proteins influenced the subcellular localization of their cognate NF-κB proteins in vertebrate cells, and their ankyrin repeat domains were predicted to interact with *Ch*-NF-κB in a manner similar to mammalian IκB and NF-κB. RNA-sequencing data from *Ch* animals indicate that *Ch*-NF-κB is expressed at high levels in early developmental stages, when *Ch*-IκB expression is low, suggesting that active *Ch*-NF-κB controls an early developmental process. In contrast, in adult animals the expression of *Ch*-IκB is high, suggesting that *Ch*-NF-κB requires a signal in order to become active. Overall, these results provide comparative information on the structure, activity, and mRNA expression of jellyfish NF-κB pathway proteins among jellyfish, and they suggest roles for NF-κB in developing and adult jellyfish.

## Introduction

Rapid increases in the number of sequenced genomes have furthered our understanding of the conservation of many genes and molecular pathways among eukaryotes, from single cells to mammals. The phylum Cnidaria occupies a key early diverging branch of the animal kingdom, and comprises approximately 11,000 extant, mostly marine organisms, including jellyfish, corals, anemones, and hydras (Steele et al., 2011). All cnidarians contain a specialized cell known as the cnidocyte, which is the phylum-specific stinging cell that is used for prey capture and defense (Babonis and Martindale, 2014).

Pattern recognition receptors (PRRs) recognize molecular structures associated with pathogens to initiate innate immune responses across metazoans. In many cases, engagement of these PRRs leads to activation of transcription factor NF-κB, which has been highly studied in mammals because of its prominent role in the regulation of immunity (Gilmore and Siggers, 2023). Emery et al. (2021) surveyed databases among various cnidarians and found that their PRRs include Toll-like receptors (TLRs), NOD-like receptors (NLRs), Retinoic acid-inducible gene I-like receptors (RLRs), and C-type lectins. Nevertheless, the types and numbers of PRRs vary considerably across cnidarians, with all four subtypes of PRRs being present in anthozoans (e.g., corals and anemones), but RLRs and C-type lectins are generally the only PRRs present in medusozoans (e.g., jellyfish and hydras). That being said, NF-κB homologs are present among all cnidarians, suggesting that downstream transcriptional networks are conserved across cnidarians (Williams and Gilmore, 2020). However, the genes and biological processes regulated by NF-κB are much less characterized in more basal organisms than in flies and mammals.

Several lines of evidence suggest that NF-κB is involved in both immunity and early life history stages in some cnidarians, including anemones and corals (Wolenski et al., 2013; Williams et al., 2018; Margolis et al., 2021). However, NF-κB has not been extensively characterized in jellyfish and hydras. Jellyfish are the free-swimming life history stage of certain gelatinous bodied members of the phylum Cnidaria. The small hydrozoan jellyfish *Clytia hemisphaerica* (*Ch*) and the moon jellyfish *Aurelia aurita* (*Aa*) have emerged as cnidarian models due to their sequenced genomes (Gold et al., 2018; Leclère et al., 2019) and, in the case of *Ch*, its small size and ability to be maintained and genetically manipulated in the laboratory (Cunningham et al., 2024).

Proteins in the NF-κB superfamily are defined by the Rel Homology Domain (RHD), which is required for dimerization, DNA binding, and nuclear translocation (Gilmore, 2006). The two subfamilies of NF-κB proteins comprise the traditional NF-κBs (p52/p100, p50/p105, *Drosophila* Relish), and the Rel proteins (RelA, RelB, c-Rel, and *Drosophila* Dif and Dorsal). NF-κB and Rel subfamily proteins differ in the types of sequences that are C-terminal to the RHD. That is, mammalian NF-κB proteins contain C-terminal inhibitory sequences consisting of a series of ankyrin (ANK) repeats, whereas Rel proteins contain C-terminal transactivation domains. Overall, the amino acid (aa) sequence of the NF-κB proteins that have been identified in cnidarians (Gilmore and Wolenski, 2012; Mansfield et al., 2017; Williams et al., 2018; Wolenski et al., 2011), poriferans (Gauthier and Degnan, 2008; Williams et al., 2020), the protist *C. owczarzaki* (Williams et al., 2021), and some choanoflagellates (Richter et al., 2018; Williams et al., 2021) more closely resemble the NF-κB subfamily of proteins (rather than Rel proteins) found in flies and vertebrates. Some basal NF-κB proteins (e.g., in the sponge *Amphimedon queenslandica*, the protist *Capsaspora owczarzaki*, and the sea anemone *Exaiptasia pallida* [*Aiptasia*]) have the RHD-ANK bipartite domain structure (like human p100 and p105), whereas others (e.g., in the anemone *N. vectensis*, *Hydra vulgaris*, and certain choanoflagellates) consist primarily of the RHD sequences, and the interacting ANK repeat sequences are encoded by separate genes, presumably due to a gene splitting event (Williams and Gilmore, 2020).

Because of the conservation of NF-κB as a likely downstream player in cnidarian immunity, we have used bioinformatic, cellular, and molecular approaches to characterize the structure, activity, regulation, and developmental expression of two jellyfish NF-κB proteins, as well as their interacting IκB-like proteins. These results indicate that although these jellyfish NF-κB proteins have many of the same properties as other metazoan NF-κB proteins, they also have sequences and activities that are distinct from NF-κB proteins of other early branching metazoans. Furthermore, we have performed RNA-sequencing analysis for gene expression of NF-κB and its interacting proteins through various developmental stages and in single cell preparations of *C. hemisphaerica*.

## 2. Materials and Methods

### 2.1. Phylogenetic analyses

The RHD sequences of NF-κB from various organisms are listed in Supplemental Table 1. Conserved RHD motifs from MEME analysis were identified based on motif predictions by Clustal Omega (Sievers et al., 2001), and each sequence was trimmed to include only the analogous RHD aa sequences. These sequences were then aligned using the MUSCLE, a phylogeny was created by PmyML, and the tree was rendered by TreeDyn as part of the “One Click” mode features from https://www.phylogeny.fr/simple_phylogeny.cgi (Dereeper et al., 2008) (Fig. 1B).

**Fig. 1.**
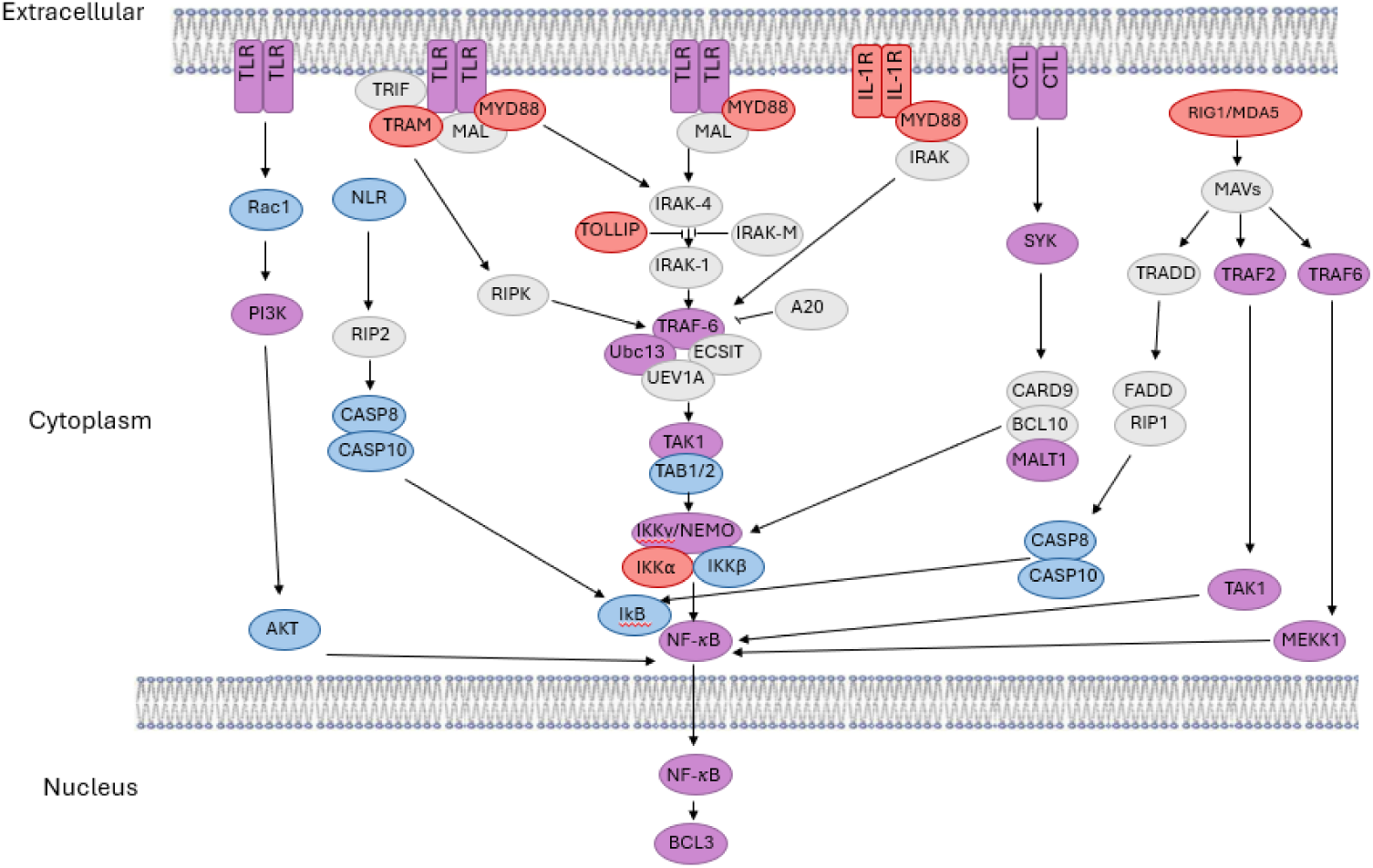
Homologs of proteins in mammalian innate immune pathways upstream of NF-κB are present in *Aurelia aurita* and *Clytia hemisphaerica*. TLR, NLR, CTL, and RIGMDA5 signaling components that are only found in humans are in grey. Homologs to human pathway components that are only in *A. aurita* are in red, homologs only in *C. hemisphaerica* are in blue, and homologs present in both are in purple. See Supplemental Table 3 for links to sequences of these jellyfish homologs.

### 2.2. Plasmid constructions

Expression plasmids for FLAG-tagged *Nv*-NF-κB and the empty pcDNA-FLAG vector have been described previously (Wolenski et al., 2011; Mansfield et al., 2017; Williams et al., 2018). The cDNA sequences of human codon-optimized *Aa*- and *Ch*-NF-κB were subcloned into pcDNA-FLAG or the yeast GAL4-fusion protein vector pGBT9. pcDNA-MYC expression vectors for the human codon-optimized *Aa* and *Ch* IκB and BCL3 proteins were created by conventional restriction enzyme-based subcloning. Details about sequences and plasmid constructions can be found in Supplemental Fig. 1 and Supplemental Table 2, respectively.

### 2.3. Cell culture and transfection

DF-1 chicken fibroblasts and human HEK 293 or 293T cells were grown in Dulbecco’s modified Eagle’s Medium (DMEM) (Invitrogen) supplemented with 10% fetal bovine serum (Biologos), 50 units/ml penicillin, and 50 μg/ml streptomycin as described previously (Wolenski et al., 2011: Williams et al., 2021). Transfection of these cells with expression plasmids was performed using polyethylenimine (PEI) (Polysciences, Inc.) as described previously (Williams et al., 2021). Briefly, on the day of transfection, cells in DMEM/10% FBS were incubated with plasmid DNA and PEI at a DNA:PEI ratio of 1:6. The media was changed ∼15-20 h after transfection, and whole-cell lysates were prepared ∼24-30 h later in AT Lysis Buffer (20 mM HEPES, pH 7.9, 150 mM NaCl, 1 mM EDTA, 1 mM EGTA, 20% w/v glycerol, 1% w/v Triton X-100, 20 mM NaF, 1 mM Na4P2O7·10H2O, 1 mM dithiothreitol, 1 mM phenylmethylsulfonyl fluoride, 1 μg/ml leupeptin, 1 μg/ml pepstatin A, 10 μg/ml aprotinin). For indirect immunofluorescence, DF-1 cells were passaged onto glass coverslips two days after transfection, were fixed with methanol, and stained the next day (Wolenski et al., 2011; Williams et al., 2021).

### 2.4. Western blotting, DNA-binding assays, reporter gene assays, and indirect immunofluorescence

Western blotting was performed using lysates prepared in AT buffer as described previously (Wolenski et al., 2011; Mansfield et al., 2017). Briefly, cell extracts were first separated on 10% SDS-polyacrylamide gels. Proteins were then transferred to nitrocellulose at 4°C at 250 mA for 2 h followed by 160 mA overnight. The membrane was blocked in TBST (10 mM Tris-HCl [pH 7.4], 150 mM NaCl, 0.1% v/v Tween 20) containing 5% powered milk (Carnation) for 1 h at room temperature. Filters were incubated at 4 °C with primary antiserum diluted in 5% milk TBST as follows: FLAG rabbit antiserum (1:1000, Cell Signaling Technology #2368) or MYC mouse antiserum (1:1000, Santa Cruz Biotechnology, #9E10). After extensive washing in TBST, filters were incubated with anti-rabbit horseradish peroxidase-linked secondary antibody (1:4000, Cell Signaling Technology) or anti-mouse horseradish peroxidase-linked secondary antibody (1:3000, Cell Signaling Technology). Immunoreactive proteins were detected with SuperSignal West Dura Extended Duration Substrate (Pierce) after exposure on a Sapphire Biomolecular Imager (Azure Biosystems).

The DNA-binding activity of NF-κB proteins was measured using a procedure that was modified from the TransAM p52 DNA-binding kit (Active Motif, #48196). Cell extracts from transfected 60-mm plates were prepared in 160 μl of AT buffer, as described above. Five μl of extract was then processed as recommended by the manufacturer, except anti-FLAG antiserum (1:1000, Cell

Signaling Technology #2368) was used as the primary antiserum. Competitions were performed using an excess of the wild-type NF-κB oligonucleotide or the mutant oligonucleotide supplied with the Active Motif kit. When the reactions started to turn blue, they were stopped, and then read at 450 nm. For each value of a reaction using a cell extract, the value obtained with AT buffer alone was first subtracted.

Yeast GAL4-site *LacZ* and 293 cell κB-site luciferase reporter gene assays were performed as described previously (Wolenski et al., 2011). Transfection and indirect immunofluorescence in DF-1 cells were performed on methanol-fixed cells that were probed with rabbit anti-FLAG primary antiserum (1:100, Cell Signaling Technology #2368), mouse anti-MYC (1:100, Santa Cruz Biotechnology, #9E10), and Alexa Fluor 488-conjugated goat anti-rabbit IgG or Alexa Fluor-555 anti-mouse secondary antiserum (Invitrogen, #A21422 ThermoFisher), as described previously (Wolenski et al., 2011; Williams et al., 2021).

### 2.5. mRNA-seq analysis of transcripts from staged C. hemisphaerica

Bulk RNA-seq data from staged *C. hemisphaerica* generated in Leclère et al. (2019) and scRNA-seq data generated in Chari et al. (2021) were reanalyzed here. Briefly, to generate bulk RNA-seq data, in Leclère et al. (2019) used male and female medusa and polyps, as well as embryos. Animals were starved for at least 24 h, kept in artificial seawater with penicillin and streptomycin, then lysed and stored at −80 °C until RNA purification. Total RNA was prepared using the RNAqueous Microkit or RNAqueous (Ambion). Purification of mRNA and construction of cDNA libraries were performed using the Kapa RNA library prep kit, and sequencing was performed using either HiSeq 2500 (single-read 50 cycles) or NextSeq (single-read 75 cycles).

RNA-seq data were collected from the following: early gastrula; 1-, 2- and 3-day-old planula; stolon, polyp head, gonozooid, baby medusa, mature medusa, male medusa (Leclère et al., 2019), growing oocyte, and fully grown oocyte (Artigas et al., 2018). RNA-seq reads were mapped using TopHat2. Genes were predicted from using Cufflinks and Cuffmerge. Herein, bulk sequencing data were accessed from the Marine Invertebrate Models Database (MARIMBA) for each gene of interest.

*C. hemisphaerica* orthologs were identified by a BLAST search of the *C. hemisphaerica* transcriptome v1.0 on MARIMBA. The gene and transcript identifiers were NF-κB: XLOC_003373, TCONS_00005923; IκB: XLOC_026731, TCONS_00042973; BCL3: XLOC_029415, TCONS_00046043

As described above, the *C. hemisphaerica* medusa single cell data generated in Chari et al. (2021) were reanalyzed to examine expression of genes of interest. In Chari et al. (2021), the scRNA-seq data were aligned to a transcriptome generated from bulk RNA-seq data. Briefly, Trinity de novo assembler was used to generate a transcriptome, and the Cufflinks Cuffcompare utility was used to merge the Trinity-assembled transcripts with any XLOC annotations from the MARIMBA v.1 transcriptome assembly.

Demultiplexing and initial processing were performed in the 10X Cell Ranger pipeline using Cell Ranger 3.0. Herein, additional analysis was done using Scanpy and kallisto bustools. Code availability: All code used to perform the analyses and generate the results and figures are available in Google Colab notebooks archived with Zenodo at https://zenodo.org/record/5519756#.YUonytNKgUE and are directly available at https://github.com/pachterlab/CWGFLHGCCHAP_2021 (from Chari et al., 2021).

## 3. Results

### Innate immune pathways in A. aurita and C. hemisphaerica

To gain an overview of possible innate immunity pathways in *A. aurita* and *C. hemisphaerica*, we used the previous analysis done by Emery et al. (2021) along with our own analysis of transcriptomic and genomic data from these two jellyfish species. As shown in Fig. 1 (and listed in Supplemental Table 3), both jellyfish have homologs of several components of the four major innate immune pathways, but generally, their pathways are less complex. Of course, we cannot rule out that our analyses missed some homologs that are present in jellyfish. One difference from previous analyses (Emery et al., 2021) was that we did find possible homologs of TLRs, but as in some other cnidarians (Brennan and Gilmore, 2018), there was no full-length plasma membrane-spanning TLR, but rather a receptor domain Leucine-rich region (LRR) protein and an internal TIR domain. The common downstream effector of these innate immune pathways, transcription factor NF-κB, was present in both of these jellyfish.

### Domain structure and phylogeny of the A. aurita and C. hemisphaerica NF-κB proteins

Our interrogation of transcriptomic and genomic databases of the jellyfish *A. aurita* and *C. hemisphaerica* identified a single NF-κB-like protein in each. Both NF-κB proteins encoded by these jellyfish consist of approximately 450 aa, containing an intact DNA-binding/dimerization Rel homology domain (RHD), but they both lacked the extended C-terminal ANK repeat domains found in mammalian NF-κB proteins (Fig. 2A).

**Fig. 2.**
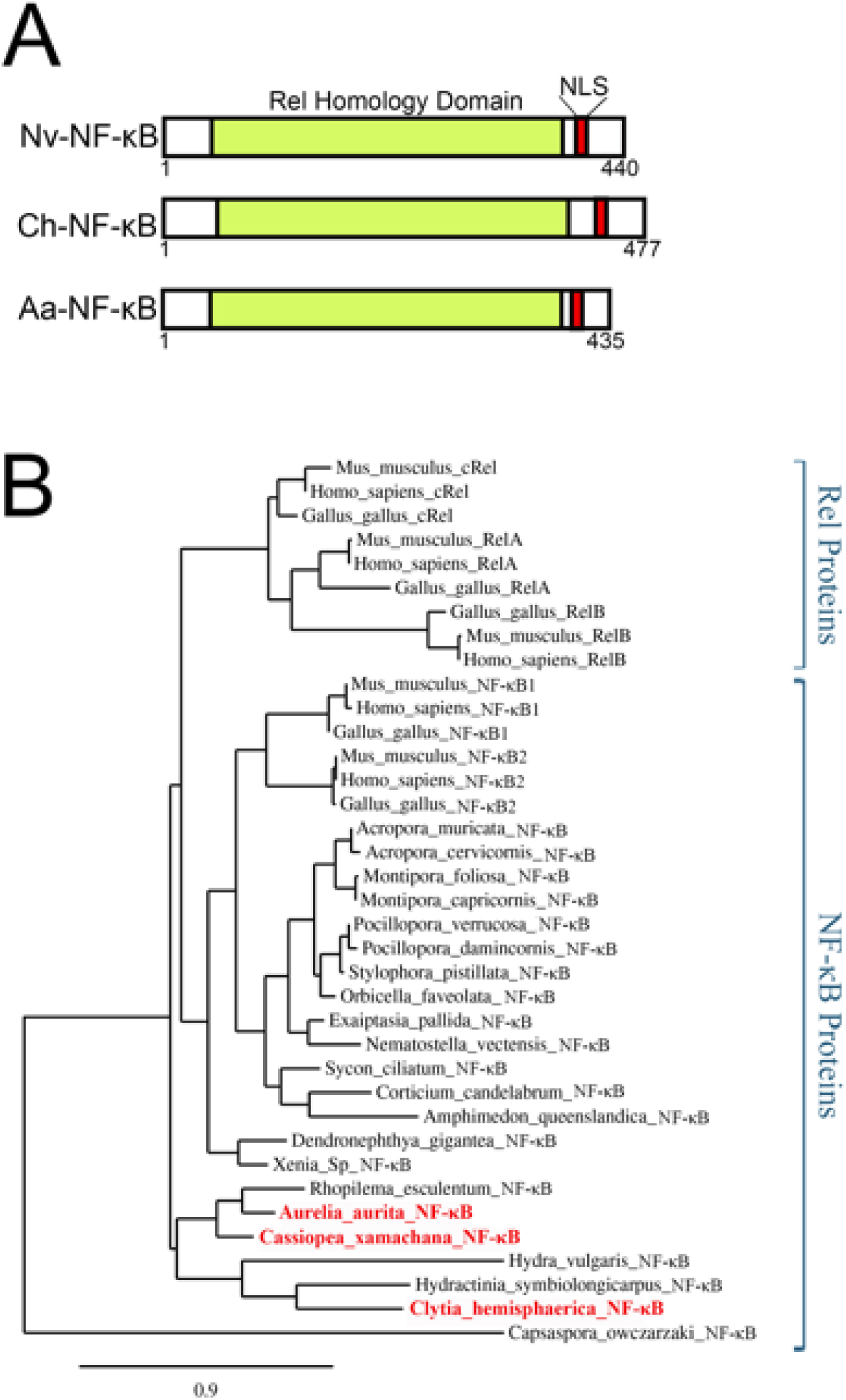
Phylogenetic and structural comparison of the *Ch*- and *Aa*-NF-κB proteins to other metazoan NF-κB proteins. (A) Schematic of jellyfish NF-κB domains: Rel Homology Domain (RHD, color), Nuclear Localization Sequence (NLS). Amino acid positions are the numbers that are below the figures. (B) Phylogenetic analysis of RHD sequences of the indicated NF-κB proteins was performed using Maximum Likelihood analysis. The phylogeny was rooted with the predicted RHD of the single-celled protist *C. owczarzaki.* The bar at the bottom of the tree indicates the scale of genetic changes as indicated by amino acid substitutions per site.

*Aa*-NF-κB is a 435 aa protein, *Ch*-NF-κB is 477 aa, and the sea anemone *N. vectensis* NF-κB is 440 aa (Fig. 2A; Supplemental Fig. 1A). A direct comparison of the two jellyfish proteins shows that the difference in the sizes of the two jellyfish NF-κB is primarily due to an additional 41 aa that are C-terminal to the NLS at the end of the RHD in *Ch*-NF-κB (Supplemental Fig. 2).

An aa sequence-based phylogenetic comparison of the RHD sequences of a variety of NF-κB proteins (Fig. 2B) showed that the RHD sequences of *Aa*- and *Ch*-NF-κB were in the large branch containing vertebrate (chicken, mouse, human) and basal NF-κB proteins, and this branch was separate from the Rel proteins defined by the vertebrate proteins (cRel, RelA, RelB). Among the basal NF-κB proteins, there were separate subclusters that contained sponge, sea anemone, and coral NF-κB proteins. Relevant to this study, the NF-κB proteins from three scyphozoan jellyfish (*Aa*, *C. xamachana, Rhopilema esculentum*) were clustered together, whereas the hydrozoan *Ch*- NF-κB protein was in a separate branch clustering with the *Hydracyntina symbiolongicarpus* and *H. vulgaris* NF-κB proteins.

### Functional characterization of jellyfish NF-κB proteins

To analyze functional aspects of these jellyfish NF-κB proteins we created plasmid expression vectors for FLAG-tagged, human codon-optimized versions of each protein (Fig. 3A). As a positive control for our experiments, we used a FLAG-tagged expression vector for the sea anemone *N. vectensis* NF-κB protein, which we have characterized previously (Wolenski et al., 2011). Anti-FLAG Western blotting of cell lysates from HEK 293T cells transfected with these vectors showed that proteins of the expected sizes were expressed in these cells, and that no such protein was expressed in cells transfected with the empty FLAG vector (Fig. 3B).

**Fig. 3.**
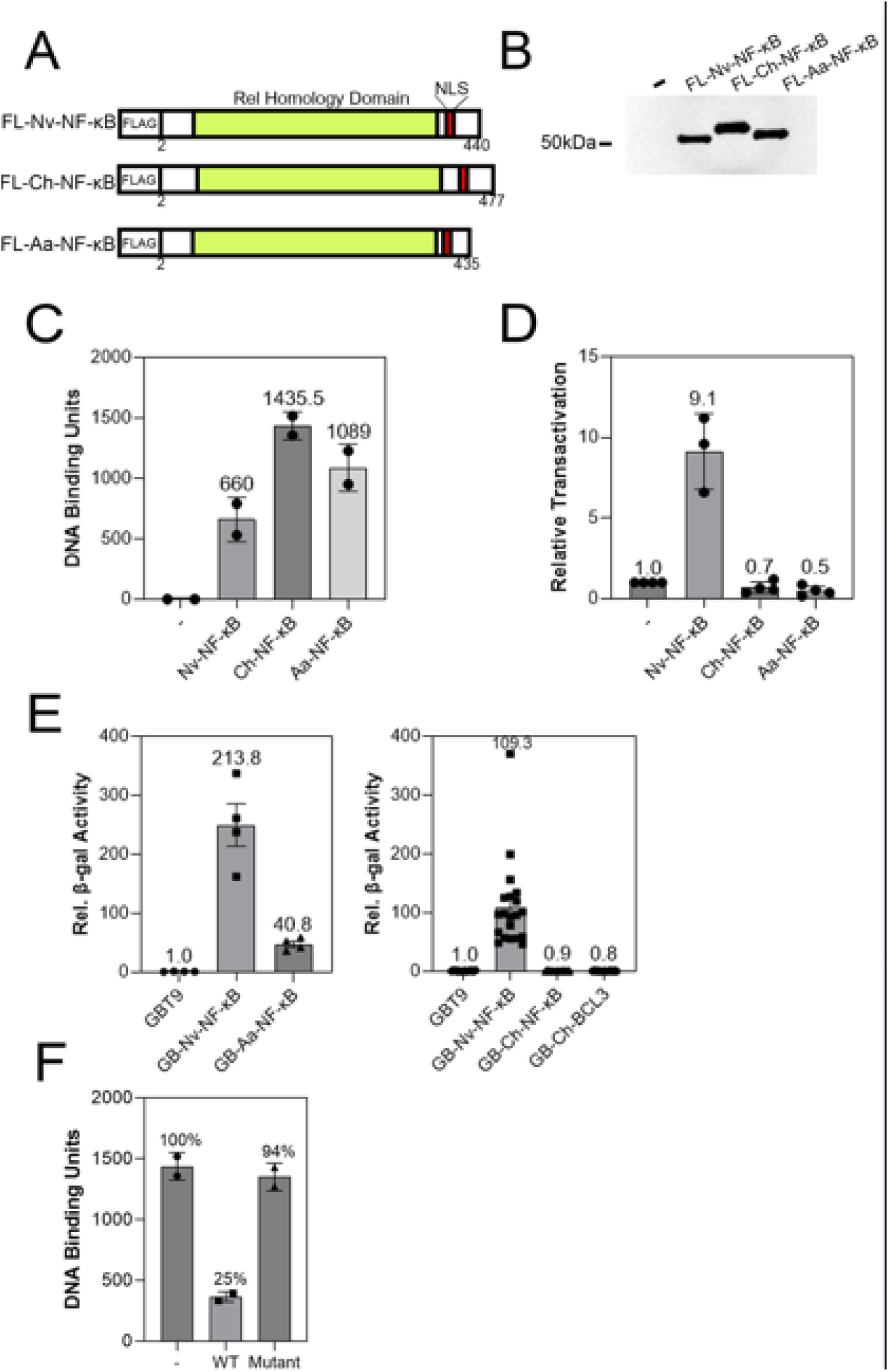
Jellyfish NF-κB proteins can bind to a mammalian DNA target site and *Aa*-NF-κB can activate transcription. (A) Depiction of the FLAG-tagged expression vectors of the *Nv*-NF-κB, *Ch*-NF-κB and *Aa*-NF-κB proteins. (B) Anti-FLAG Western blot of 293T cell lysates transfected with the indicated expression vectors. (C) κB site DNA-binding activity from 293T cell extracts (in B) that were transfected with the indicated FLAG-tagged proteins. Shown are the results of two independent experiments, with the relative DNA-binding units of the ELISA-based DNA-binding assays (see Materials and Methods). (D) A κB-site luciferase reporter gene assay was performed with the indicated proteins in 293 cells. Luciferase activity is relative (Rel.) to that seen with the empty vector control (1.0), and values are the averages of three assays performed with triplicate samples with standard error. (E) An ELISA-based DNA-binding assay was performed as in (C) except an excess of wild-type competitive DNA (WT) or a mutant oligonucleotide (Mutant) was added before adding the extracts. A control assay with no competing DNA (-) was also performed, and was adjusted to 100%. (F) A GAL4-site *LacZ* reporter gene assays were performed in yeast Y190 cells. β-galactosidase (β-gal) reporter gene activity is relative (Rel.) to the GAL4 (aa 1-147) control (1.0). Values are averages of two assays performed with duplicate samples with standard error.

To analyze the DNA-binding ability of these proteins, extracts from these transfected cells were subjected to a modified indirect ELISA-based assay in a 96-well plastic plate that contained an NF-κB-site oligonucleotide bound to each well. Proteins from the extract that bound to this oligonucleotide were then detected with anti-FLAG antiserum and a horseradish peroxidase-conjugated secondary antibody, which was used for enzymatic quantification of DNA-bound proteins. As shown in Fig. 3C, extracts from cells transfected with the jellyfish NF-κB protein vectors contained over 1000-fold more NF-κB-site oligonucleotide binding activity than the low level seen in extracts from the empty vector-transfected negative control cells. Overall, both jellyfish NF-κB proteins have NF-κB-site DNA-binding activity that is comparable to the DNA-binding activity of extracts from the *Nv*-NF-κB-expressing positive control cells (660-fold increased DNA-binding activity).

We next analyzed the ability of the two jellyfish NF-κB proteins to activate transcription from an NF-κB-site luciferase reporter gene when transfected into HEK 293 cells (Fig. 3D). Consistent with our previous results, *Nv*-NF-κB activated transcription approximately 10-fold over what was seen in empty vector-transfected cells. In contrast, neither *Aa*-NF-κB nor *Ch*-NF-κB showed an ability to activate this NF-κB-site reporter gene in these assays.

We also determined the ability of these NF-κB proteins to activate transcription as GAL4-fusion proteins in yeast, using a yeast strain with an integrated *lacZ* reporter locus with five upstream GAL4 binding sites. We have previously used such assays to measure the inherent ability of basal NF-κB proteins to activate transcription, in part because yeast do not have NF-κB proteins and in part because these assays assess the ability of the GAL4-fusion protein to activate the reporter gene but do not rely on the ability of the NF-κB protein to bind to DNA by itself. As shown in Fig. 3E, GAL4-*Aa*-NF-κB showed over 40-fold greater reporter gene β-galactosidase activity as compared to yeast cells expressing only the GAL4 DNA-binding domain (aa 1-147). In contrast, GAL4-*Ch*-NF-κB did not increase β-galactosidase activity as compared to the GAL4 aa 1-147 control. As shown previously (Wolenski et al., 2011), GAL4-*Nv*-NF-κB robustly activated transcription (over 100-fold) in yeast. Of note, a GAL4-fusion protein containing a predicted *Ch*-BCL3 protein, which can act as a co-activator for NF-κB in mammals, did not activate transcription.

Taken together, the results in this section indicate that *Aa*- and *Ch*-NF-κB can both bind to a consensus NF-κB site, however, neither NF-κB protein can activate transcription in human cells from a reporter gene having upstream NF-κB binding sites. To rule out the possibility that the NF- κB-site DNA-binding activity of the jellyfish NF-κB proteins was due to non-specific DNA-binding activity, we performed a competition assay in the ELISA-based DNA-binding assay. As shown in Fig. 3F, a wild-type NF-κB oligonucleotide competed for the high level of DNA-binding activity seen with the lysates containing *Ch*-NF-κB, whereas a mutant oligonucleotide did not compete for the *Ch*-NF-κB DNA-binding activity. Therefore, the ability of *Ch*-NF-κB to bind to DNA is specific for the presence of an NF-κB site in this assay.

### Structural predictions of the jellyfish NF-κB proteins

Based on their sequence and DNA-binding similarities, we next used AlphaFold3 (https://alphafoldserver.com; Abramson et al., 2024) to generate predicted structures of *Ch*- and *Aa*-NF-κB dimers and, as control, the mouse NF-κB p50 dimer, all bound to a consensus NF-κB site (GGGAATTCCC). As shown in Fig. 4, the predicted structures of all three NF-κB proteins are quite similar: that is, with clear DNA-binding and dimerization regions and with a central area that encircles the target DNA.

**Fig. 4.**
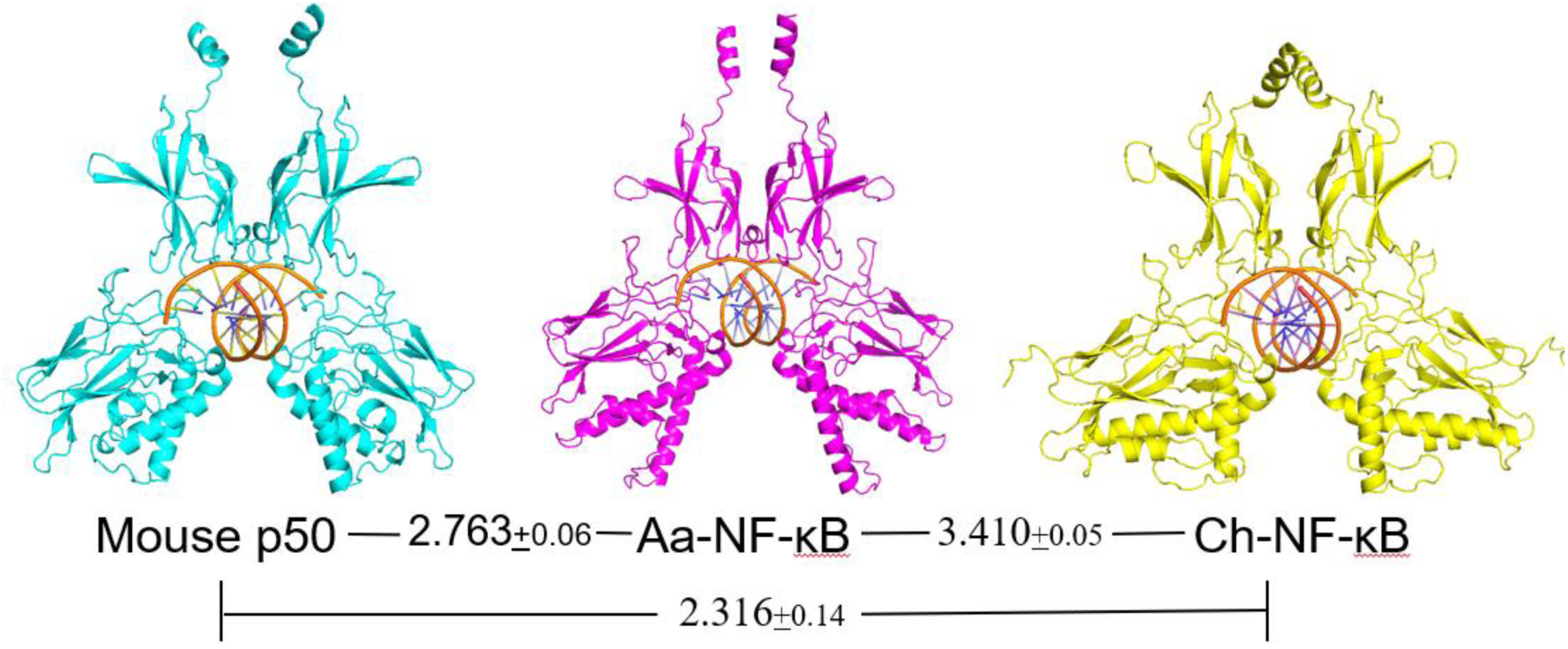
Alphafold3-based structures of jellyfish *Ch*- and *Aa*-NF-κB proteins on a palindromic NF-κB binding site (TGGGAATTCC). Double-stranded DNA is in orange. Values are the average RMSD values (plus standard deviation) for three independent comparisons of the indicated AlphaFold3-generated structures.

To more quantitatively compare the jellyfish NF-κB dimers to each other and to mammalian NF- κB p50, we exported the AlphaFold3 structures into PyMOL (www.pymol.org) and then compared the various dimers as they would appear on bound DNA, using Root Square Mean Deviation (RMSD) values as a comparison factor. As a control, we first compared the predicted mouse NF-κB p50 dimer to the X-ray crystal structure-derived mouse p50 structure (Ghosh et al., 1995); this comparison gave a low RMSD (0.831±0.17), indicating that the AlphaFold3 prediction is in good agreement with the experimentally derived structure for mouse p50 on DNA. Surprisingly, the two jellyfish NF-κB dimers were slightly less similar to each other (RMSD, 3.410±0.05) than each NF-κB dimer was to mouse p50 NF-κB (*Aa*-NF-κB è mouse p50, RMSD 2.763±0.06 and *Ch*-NF-κBè mouse p50 RMSD 2.316±0.14).

### Interaction of jellyfish NF-κB and IκB proteins

Active NF-κB proteins are generally located in the nucleus. To assess the subcellular localization of the *Aa*- and *Ch*-NF-κB proteins we transfected chicken DF-1 cells with our FLAG-tagged jellyfish NF-κB expression vectors, and then performed anti-FLAG indirect immunofluorescence. DF-1 cells were used because they show a spread-out morphology that is useful for such subcellular localization studies. As shown in Fig. 5A and Table 1, *Ch*-NF-κB and *Aa*-NF-κB, as well as the control *Nv*-NF-κB (Wolenski et al., 2011), were located either totally or partially in the nucleus in over 90% of the transfected DF-1 cells.

**Fig. 5.**
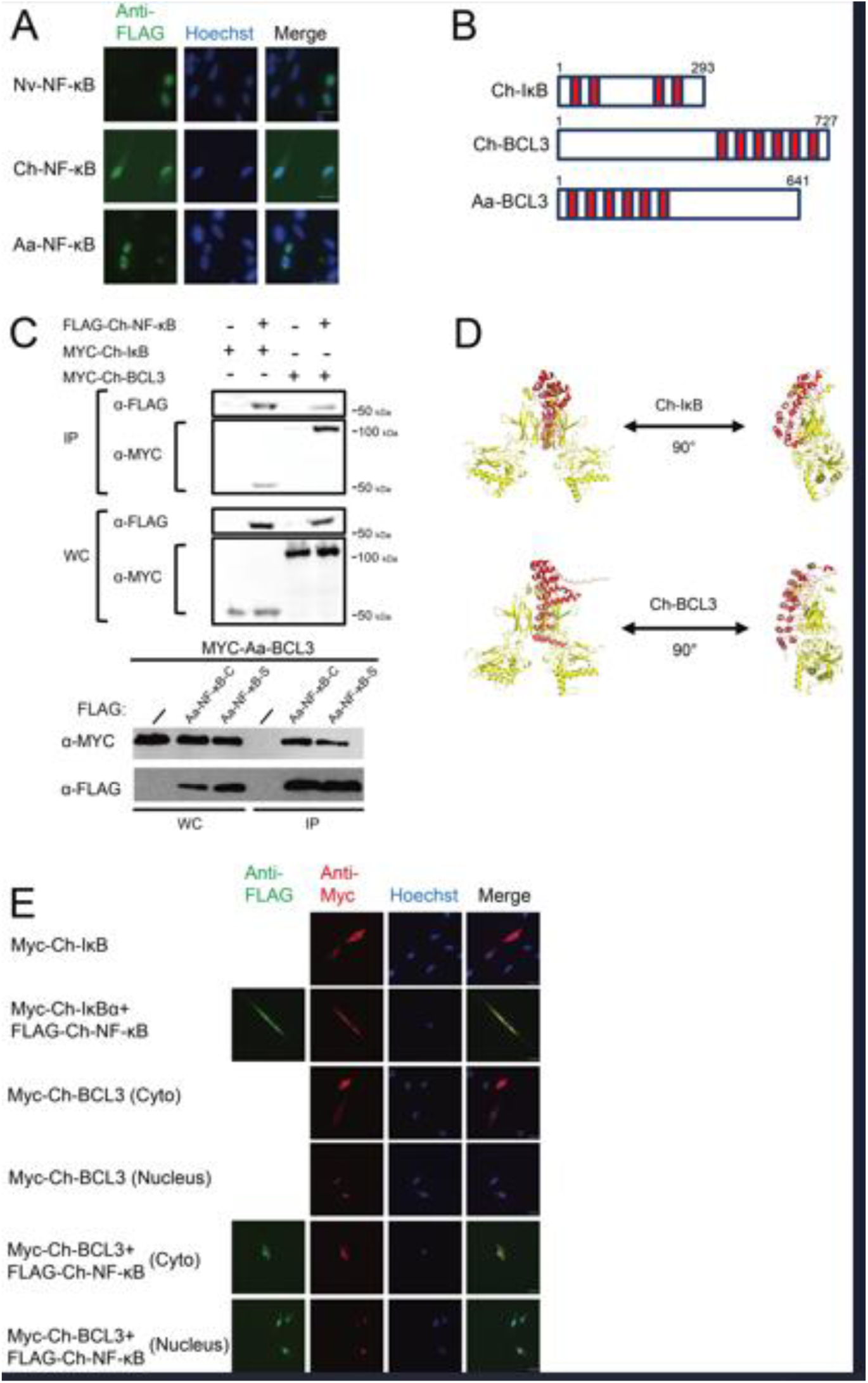
Jellyfish IκB and BCL-3 proteins can bind their respective NF-κB proteins and dictate subcellular localization. (A) Nuclear localization of *Nv*-, *Ch*-, and *Aa*-NF-κB proteins in transfected DF-1 cells. Green, anti-FLAG immunofluorescence; Blue, Hoechst nuclear staining. (B) Structures of the IκB and BCL3 proteins used in these assays. (C) Co-immunoprecipitation of the indicated proteins from HEK A293T-transfected cell extracts. Anti-FLAG immunoprecipitations were performed on extracts from cells transfected with the indicated plasmids. Next, anti-FLAG or Anti-MYC Western blotting was performed. Input represents 7.5% of the lysates used in the corresponding anti-FLAG pulldown experiments. (D) Alphafold3-based structures of *Ch*-NF-κB (yellow) with the ANK repeat domains of *Ch*-BCL3 and *Ch*-IκB. (E) DF-1 chicken fibroblasts were transfected with the indicated pcDNA expression vectors, and two days later cells subjected to anti-FLAG and/or anti-MYC indirect immunofluorescence. Anti-FLAG, green; anti-MYC; Hoechst-stained nuclei, blue; and merged images are shown. Quantitation of cell staining is reported in Table 1.

**Table 1.** Subcellular localization of over-expressed jellyfish NF-κB pathway proteins in chicken DF-1 cells.

| **Protein** | **%Nuclear** | **%N/C** | **%Cytoplasm** | **%N/N** | **%C/C** |
| --- | --- | --- | --- | --- | --- |
| **Nv-NF-κB** | 87.3 (511) | 8.9 (52) | 3.8 (22) |  |  |
| **Ch-NF-κB** | 86.9 (158) | 4.9 (9) | 8.2 (15) |  |  |
| **Ch-IκB** |  | 6.3 (9) | 93.7 (133) |  |  |
| **Ch-BCL3** | 67 (218) | 10.1 (33) | 23 (75) |  |  |
| **Ch-NF-κB + IκB** |  |  |  |  | 100 (27) |
| **Ch-NF-κB + BCL3** |  |  |  | 80 (80) | 20 (20) |

| **Protein** | **%Nuclear** | **%N/C** | **%C** | **%N/N** | **%NC/NC** | **%C/C** |
| --- | --- | --- | --- | --- | --- | --- |
| **Aa-NF-κB** | 70.3 (409) | 26.3 (153) | 3.4 (20) |  |  |  |
| **Aa-BCL3** | 56.5 (288) | 34.1 (174) | 9.4 (48) |  |  |  |
| **Aa-NF-κB + BCL3** |  |  |  | 67.8 (335) | 15.4 (76) | 16.8 (83) |

Numbers in parentheses are the number of cells counted for the given protein localization.

Mammalian NF-κB proteins are well-known for their ability to interact with IκB family proteins (Morin and Gilmore, 1993), which are proteins that contain multiple ANK repeat sequences that bind to RHD sequences of NF-κB and Rel proteins. A search of *Ch* genomic and transcriptomic sequence databases identified single transcripts encoding putative IκB and BCL3 proteins. The general structures of these proteins and their sequences, with their designated ANK repeats, are shown in Fig. 5B and Supplemental Fig. 1.

To determine whether these IκB-like proteins could interact with *Ch*-NF-κB, we first created MYC-tagged expression vectors for the *Ch*-IκB and *Ch*-BCL3 proteins. We then co-transfected HEK 293T cells with expression vectors for the FLAG-tagged *Ch*-NF-κB and the MYC-tagged *Ch*-IκB or *Ch*-BCL3, performed anti-FLAG bead immunoprecipitations, and probed for FLAG-NF-κB and either MYC-IκB or MYC-BCL-3. As shown in Fig. 5C, FLAG-*Ch*-NF-κB pulled down both *Ch*-IκB and *Ch*-BCL3 in these experiments, which were not pulled down in cells transfected with the FLAG vector alone. Moreover, using AlphaFold3, the ANK repeat regions of *Ch*-IκB and *Ch*-BCL3 each docked on the *Ch*-NF-κB protein (Fig. 5D) in a manner that was consistent with an ANK repeat-RHD interaction that has been shown to occur by X-ray crystallography with mammalian IκBs on NF-κBs (Huxford et al., 1998). Furthermore, the ANK repeat structures are consistent with what one would predict for a protein with 4-6 ANK repeats (Kane and Spratt, 2021).

We next looked at the effect of the *Ch*-IκB and *Ch*-BCL-3 proteins on the subcellular localization of Ch-NF-κB in DF-1 cells by indirect immunofluorescence. As shown in Fig. 5E, *Ch*-IκB was located exclusively in the cytoplasm of over 90% of transfected cells when expressed alone. When *Ch*-IκB and *Ch*-NF-κB were co-expressed, both proteins were primarily located in the cytoplasm, which is consistent with the ability of *Ch*-IκB to retain NF-κB in the cytoplasm in other mammalian and non-mammalian systems (Morin and Gilmore, 1993).

BCL3 has been shown to be both a cytoplasmic retention protein and a nuclear co-activator for NF-κB transcription factors (Liu et al., 2022; Wolenski et al., 2011). When expressed alone, Ch-BCL3 showed heterogeneous subcellular localization: located in the cytoplasm (∼23% of cells), nucleus (∼67%), or both (∼10%) in different cells (Fig. 5E and Table 1). Consistent with the ability of *Ch*-BCL3 to interact with *Ch*-NF-κB (Fig. 5C), *Ch*-NF-κB and *Ch*-BCL3 showed nuclear or cytoplasmic co-localization in a given cell where they were co-expressed (Fig. 5E and Table 1). These results, together with their ability to be co-immunoprecipitated from cells (Fig. 5C), suggest that *Ch*-NF-κB and *Ch*-BCL3 are in a cytoplasmic or nuclear complex in any given cell.

For *Aa*, we identified a single IκB-like protein that (by BLAST) appeared most similar to IκBε and BCL3. For the sake of this paper, we will call this protein Aa-BCL3. In a co-immunoprecipitation experiment using transfected HEK 293T cell lysates, FLAG-Aa-NF-κB pulled down MYC-Aa-BCL3 (Fig. 5C). When transfected alone, *Aa*-BCL3 showed heterogeneous localization in DF-1 cells: nucleus (56.4%), nucleus/cytoplasm (34.2%) and cytoplasmic (9.4%). Moreover, *Aa*-BCL3 and *Aa*-NF-κB co-localized when co-expressed in DF-1 cells (see Table 1), similar to the analogous *Ch* proteins.

### Expression of NF-κB, IκB and BCL3 mRNAs in C. hemisphaerica

To gain insight into possible biological role(s) of NF-κB in *C. hemisphaerica*, we looked at the relative expression of NF-κB, BCL3 and IκB transcripts in previously created mRNA databases of *C. hemisphaerica* developmentally staged tissues (Leclère et al., 2019). From the transcript expression data shown in Fig. 6A, there are several readily apparent take-aways: 1) the NF-κB and BCL3 transcripts show their highest levels of expression in gastrula and planula stages; 2) in contrast, IκB transcript expression is lowest in gastrula and planula (when NF-κB and BCL3 transcripts are at their highest) and IκB is expressed at its highest level in gonad cells and medusa; and 3) tissues from the polyp stages show intermediate levels of expression for all three of transcripts.

**Fig. 6.**
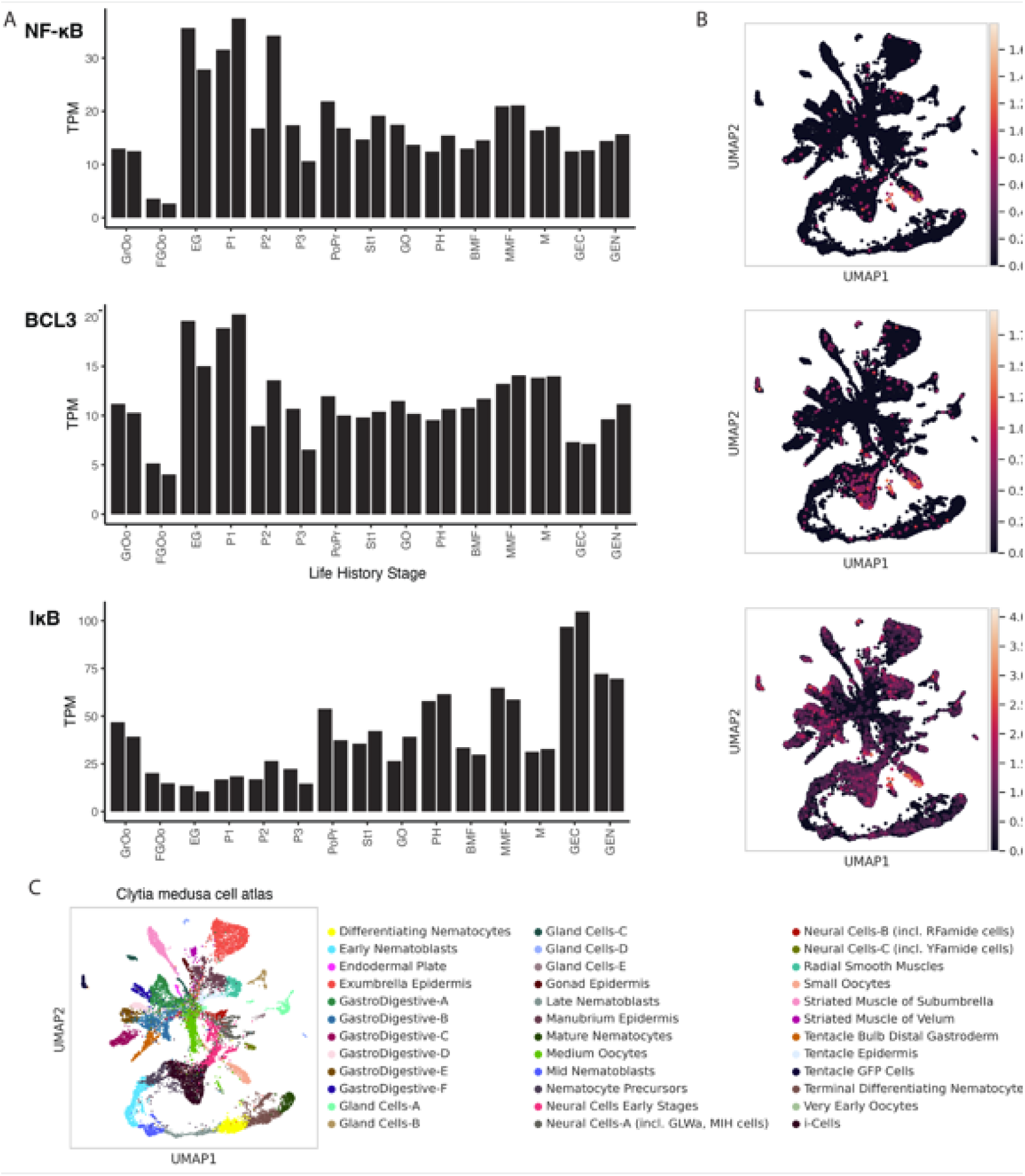
Bulk and single cell RNA-sequencing reveal expression patterns of NF-κB, BCL3 and IκB from *C. hemisphaerica.* (A) Barplots show bulk RNA sequencing data quantified as transcripts per million (TPM) for each gene and multiple life history stages. GrOo (growing oocyte), GFOo (fully grown oocyte), EG (early gastrula), P1 (planula 1), P2 (planula 2), P3 (planula 3), PoPr (primary polyp), stolon (St), GO (gonozoid), PH (polyp head), BMF (baby medusa female), MMF (mature medusa female), M (mature medusa male), GEN (gonad endoderm), GEC (gonad ectoderm). (B) Several cell types were identified in a single cell atlas of gene expression from *Clytia hemisphaerica* medusae. UMAP plots show each cluster in a different color, with cluster identities labeled based on gene expression. Cells expressing NF-κB expression were found primarily in oocytes and, to a lesser extent, in select other cell types. IκB expression was also strongly detected in oocytes, as well as I-cells, and several gastrodigestive cell clusters, among others. BCL3 was expressed mainly in oocytes and I-cells. (C) Cell Atlas shows cell types of *C. hemisphaerica* medusae, identified with different colors.

Single-cell RNA-sequencing from whole medusa revealed expression of NF-κB, BCL3 and IκB transcripts in several genetically identified cell types (Figs. 6B and 6C). Oocytes were the primary cell type that showed expression of all three NF-κB, IκB, and BCL3 mRNAs. Each of these genes were also detected at low levels in several other cell types, including neurons. The stem-cell population in cnidarians, known as I-cells, showed expression of IκB and BCL3, but not NF-κB, at this level of analysis. IκB was also notably detected in several gastrodigestive cell clusters.

## 4. Discussion

In this report, we demonstrate that two jellyfish model organisms have active NF-κB DNA-binding proteins. As such, the two jellyfish that we have studied here are similar to most evolutionarily basal organisms in that their genomes encode one NF-κB protein and no Rel proteins. Using an aa sequence-based phylogenetic analysis, the *Aa* and *Ch* NF-κBs are shown to be similar phylogenetically and structurally to other metazoan NF-κB proteins (Fig. 2B). In addition, their DNA-binding abilities, AlphaFold3-predicted structures, subcellular localizations, and interactions with IκB proteins are similar to NF-κB proteins found in several other species. Developmental mRNA expression profiling for *C. hemisphaerica* is consistent with its NF-κB having a role in early development and in activated adult immunity.

It is clear from our results (Fig. 1 and Supplementary Table 3) and those of Emery et al. (2021) that *Aa* and *Ch* have genes encoding many of the upstream components of the four standard metazoan innate immune pathways. Nevertheless, these pathways are less complex than those in humans. In contrast to Emery et al. (2021), we identified possible split TLR sequences, i.e., possible separate LRR domain and TIR domain proteins. Similar split TLR-like proteins have also been found in the sea anemone *Aiptasia* and *Hydra vulgaris* (Brennan and Gilmore, 2018). In addition, there is a MyD88 adapter protein, which binds to TIR domains for intracellular signaling from TLR-like proteins, in *Aa* that suggests that TLR-like signaling can occur in this jellyfish. Indeed, whether any of these putative pathways shown in Fig. 1 function in immune-like pathways in these two jellyfish is not known.

Assuming that the bulk mRNA expression levels from *Ch* animals (Fig. 6A) correlate with protein expression, then one might predict that the high expression of NF-κB mRNA, coupled with low levels of IκB mRNA expression, at early stages of development (gastrula and planula) result in high levels of active, nuclear NF-κB protein at these stages. In contrast, one would expect NF-κB to be primarily inactive at later life stages (i.e., in polyps and medusae) when IκB mRNA expression is high. This model is consistent with what has been shown in the sea anemone *N. vectensis*, where NF-κB is detected in the nucleus of cells at the gastrula stage (Wolenski et al., 2013), but is located in the cytoplasm of cnidocytes, bound to IκB, in mature anemones (Wolenski et al., 2013). Moreover, we have previously shown that morpholino-based knockdown of NF-κB in the sea anemone *N. vectensis* embryo prevents the later formation of mature cnidocytes (Wolenski et al., 2013). Based on these results, we speculate that nuclear NF-κB in *C. hemisphaerica* is also involved in the development of early precursor cells. This model for an early developmental role for the *Ch*-NF-κB pathway is supported by the expression of NF-κB, BCL3, and IκB in oocytes (Fig. 5B).

Based on bulk RNA expression from later life stages of *Ch* (i.e., polyps and medusae) Ch-NF-κB may be retained in the cytoplasm by high levels of IκB, and thus, the induction of nuclear NF-κB activity may require signal-induced degradation of IκB or transcriptional activation of NF-κB expression. We note that *Ch*-IκB has two serines near its N terminus (S14 and S20) that are in a similar sequence context to serine residues that are IκB kinase phosphorylation sites in human and *N. vectensis* IκB proteins and that promote proteasomal degradation of IκB when phosphorylated (Supplemental Fig. 3). In adult *Ch*, we speculate that the induction of active NF-κB activity regulates an immune response. Consistent with that hypothesis, Emery et al. (2024) have recently shown in the upside-down jellyfish *Cassiopea xamachana* that NF-κB mRNA expression is increased by exposure to a bacterial pathogen.

In the symbiotic sea anemone *Aiptasia*, NF-κB is found in the nucleus of cells even in adult tissues (Mansfield et al., 2017). In contrast, NF-κB protein is present in the cytoplasm of cells in adult *N. vectensis* anemones and, as we note above, likely also in *Ch*. There are several differences between *C. hemispherica*/*N. vectensis* and *Aiptasia*, which may account for differences in regulation and subcellular localization of their NF-κB proteins. First, the NF-κB protein in *Aiptasia* is a bipartite NF-κB wherein the ANK repeat IκB sequences are included in the C terminal half of its NF-κB (Mansfield et al., 2017), whereas *C. hemispherica* and *N. vectensis* both have separate genes encoding NF-κB and IκB (Wolenski et al., 2011). Indeed, in *Aiptasia* NF-κB is found as a constitutively processed and nuclear protein in mature anemones (Mansfield et al., 2017), whereas in *N. vectensis* NF-κB and IκB are co-expressed in the cytoplasm of an overlapping set of cells in mature anemones (Wolenski et al., 2013). Moreover, NF-κB is primarily expressed in adult cnidocytes in *N. vectensis* (Wolenski et al., 2013), but *Aiptasia* NF-κB is largely expressed in gastrodermal cells, which host algal symbionts (Mansfield et al., 2017). Finally, *N. vectensis* (and *C. hemisphaerica*) do not form any known symbiosis with algae as occurs in *Aiptasia* and many stony corals. Overall, the data to date suggest that NF-κB has different biological roles and regulation in different cnidarians, and its structure and regulation may reflect the different biological processes that NF-κB regulates in a given cnidarian species. Of note, NF-κB mRNA is suppressed in several organisms when they are hosting algal symbionts, including the anemone *Aiptasia* (Mansfield et al., 2017), the jellyfish *C. xamachana* (Emery et al., 2024), and the salamander *Ambystoma maculatum* (Burns et al., 2017).

Ankyrin repeat-containing IκB-like proteins from both *Aa* and *Ch* can interact with and influence the subcellular localization of their cognate NF-κB proteins in cell-based assays (Fig. 4, Table 1). The *Ch*-IκB protein that we have identified appears to be a standard cytoplasmic retention protein for *Ch*-NF-κB. The protein that we have designated as *Ch*-BCL3 shows both cytoplasmic and nuclear localization when expressed in vertebrate cells (Fig. 5, Table 1). We have found similar IκB and BCL3 proteins in *N. vectensis* (Wolenski et al., 2011). In mammals, BCL3 can act as both a cytoplasmic sequestering protein and a co-activator for NF-κB (Liu et al., 2022). BCL3 may perform similar roles for *Ch*-NF-κB, and we especially note that the profile of *BCL3* mRNA expression essentially parallels that of NF-κB mRNA in developing *Ch* (Fig. 5A).

Our AlphaFold3 structural predictions and comparisons of the two jellyfish NF-κB dimers on a consensus DNA have several interesting implications. The comparison of the AlphaFold3-predicted structures of the NF-κB’s on DNA (Fig. 4) indicate that *Aa*-NF-κB and *Ch*-NF-κB structures are slightly less similar to one another than either is to the mouse NF-κB p50 structure. This is in contrast to the strict aa-based phylogenetic analysis (Fig. 2B) that indicates that *Aa*-NF-κB and *Ch*-NF-κB are more similar to one another than either is to mouse NF-κB p50. Therefore, Alphafold3 comparisons may, at least in some cases, be more relevant to a protein’s activity than a strict phylogenetic aa comparison.

Using human HEK 293 cells, we have previously shown that NF-κB proteins from a variety of basal organisms—including sea anemones, corals, sponges, and protists (Wolenski et al., 2011; Mansfield et al., 2017; Williams et al., 2018; Williams et al., 2020; Williams et al., 2021)—can activate transcription of the same κB-site reporter used in the experiments described herein. However, neither jellyfish NF-κB could activate transcription of this NF-κB-site reporter locus in HEK cells, under conditions where the sea anemone *Nv*-NF-κB protein readily activated this reporter (Fig. 3D). Nevertheless, *Aa*- and *Ch*-NF-κB both robustly bound DNA when expressed in HEK cells, and they did so to an even greater extent than the *Nv*-NF-κB protein (Fig. 3C). Thus, if the *Aa*- and *Ch*-NF-κB proteins are activators of transcription in jellyfish, they likely require jellyfish-encoded proteins or modifications to activate transcription in their native environments. In this regard, the jellyfish NF-κB proteins resemble the *Drosophila melanogaster* NF-κB Relish protein that requires phosphorylation (Ertűrk-Hasdemir et al., 2009) and interaction with a specific co-activator (Cammarata-Mouchtouris et al., 2020) in order to activate certain fly immunity genes. Similar to the jellyfish NF-κB proteins we have characterized here, human NF-κB p50 and p52 homodimers do not by themselves activate κB-site genes and require interaction with BCL3 to activate transcription (Liu et al., 2020); that being said, *Ch*-BCL3 did not activate transcription as a GAL4-fusion protein in yeast.

Our results provide new information on the biochemical and molecular properties and biological roles of NF-κB proteins in basal organisms, namely jellyfish. As such, they contribute to our understanding of the diversification of this transcription factor that regulates many immune and inflammatory processes across a broad swath of modern-day phyla.

## Acknowledgments

This research was supported by National Science Foundation grant IOS-1354935 (T.D.G.). J.P and K.A. were supported by the Boston University (BU) Undergraduate Research Opportunities Program, and A.N.-R. and J.P. were supported by an NSF-REU (to T.D.G.). B.W. was supported by the Freedom Together Foundation. Designated authors performed research as part of the undergraduate molecular biology project laboratory course BB522 (Spring, 2022 and 2024), and were supported by funds from the BU Biology Department. Special thanks to Todd Blute (BU) for help with microscopy, to Michael Garbati (Active Motif) for help with the DNA-binding assay, and Maria Valadez-Ingersoll (BU) for helpful discussions.

## Supplemental Author Information

BB522 Molecular Biology Laboratory Emma Batchelor, Cameryn Boggio-Shean, Sean Bridges, Emily Budram, John Paul Capuano, Charles Courtemanche, Melisa Cruz, Simai Deng, Ava DiStefano-Forti, William Dorst, Cecilia Downs, Evangeline Earnest, Joseph Friedlander, Miguel Galera Diez, Daniel Garcia, Isabella Garza, Victoria Gauntner, Sean Gow, Mano Harada, Chae Yoon Jhang, Grace Johnson, Gabrielle Kim, Ryan Lawrence, Chun Lin, Jacy Marquez, Rhianna Murphy, Nirmiti Naik, Raja Poda, Steven Rubin, Griffin Salus, Sakshi Shah, Jaimu Sun, Abdulmalk Tahlil, Cole Varela, Megan Wong, Kalin Xu, Carmen Zhou, Timothy Zhu; Anisah Adeagbo, Christian Badawi, Kimia Bazarganpour, Evan Blade, Yujin Chen, Edmund Cheung, Yuk Hei Cheung, Sophia Cornish,Tyler Critz, Tsetan Dhondup, Bridget Dibbini, Daisy DiGregorio, Luciano Foranoce, Bianca Frintu, Paul Hee, Giovi Hersch, Julia Hetzko, Isabella Hirt, Julia Hoffman, Saiyan Joseph, Sarah Josinsky, Ethan Leviss, Jiayi Li, Kevin Li, Charmaine Liu, Madison Marano, Itthipun Munkong, Alex Novakovic, Christopher Parcels, Sophie Pastor, Anton Sergeev, Swastika Sharma, Zicheng Sun, Purnima Venkat, Karisma Verma, Xinyi Yang, Raymond Zeng, Liyuan Zhu (Programs in Biochemistry & Molecular Biology and in Biology, Boston University, Boston, MA, USA)

## Abbreviations used are

aa: amino acid(s)
Aa: Aurelia aurita
Ch: Clytia hemisphaerica
IκB: inhibitor of kappaB binding
NF-κB: nuclear factor kappaB
Nv: Nematostella vectensis
PBS: phosphate-buffered saline

## REFERENCES

Abramson., J., Adler, J., Dunger, J., Evans, R., Green, T., Pritzel, A., Ronneberger, O., Willmore, L., Ballard, A.J. , Bambrick, J., Bodenstein, S.W., Evans, D.A., Hung C.-C., O’Neill, M.O., Reiman, D., Tunyasuvunakool, K., Wu, Z., Žemgulytė, A., Arvaniti, E., Beattie, C., Bertolli, O., Bridgland, A., Cherepanov, A., Congreve, M., Cowen-Rivers, A.I., Cowie, A., Figurnov, M., Fuchs, F.B., Gladman, H., Jain, R., Khan, Y.A., Low, C.M.R., Perlin, K., Potapenko, A., Savy, S., Singh, S., Stecula, A., Thillaisundaram, A., Tong, C., Yakneen, S., Zhong, E.D.,, Zielinski, M., Žídek,, A., Bapst, V., Kohli, P., Jaderberg, M., Hassabis, D., Jumper, J.M., 2024. Accurate structure prediction of biomolecular interactions with AlphaFold3. Nature 630, 493–500.

Artigas, G.Q., Lapébie, P., Leclère, L., Takeda, N., Deguchi, R., Jékely, G., Momose, T., Houliston, E., 2018. A gonad-expressed opsin mediates light-induced spawning in the jellyfish *Clytia*. eLife 7, e29555.

Babonis, L.S., Martindale, M.Q., 2014. Old cell, new trick? Cnidocytes as a model for the evolution of novelty. Integ. Compar. Biol, 54, 714–722.

Böhm, M., Hentschel, U., Friedrich, A., Fieseler, L., Steffen, R., Gamulin, V., Müller, I., Müller, W., 2001. Molecular response of the sponge *Suberites domuncula* to bacterial infection. Mar. Biol. 139, 1037–1045.

Brennan, J.J. Gilmore, T.D. 2018. Evolutionary origins of Toll-like receptor signaling. Mol. Biol. Evol. 35, 1586–1587.

Burns, J.A., Zhang, H., Hill, E., Kim, E., Kerney, R., 2017. Transcriptome analysis illuminates the nature of the intracellular interaction in a vertebrate-algal symbiosis. Elife 6, 1–32.

Cammarata-Mouchtouris, A., Nguyen, X.-H., Acker, A., Bonnay, F., Goto, A., Fauvarque, M.-O., Boutros, M., ReiSchart, J.-M., Matt, N., 2020. Hyd ubiquitinates the NF-κB co-factor Akirin to operate an effective immune response in *Drosophila*. PLOS Pathog. 16, e1008458.

Chari, T., Weissbourd, B., Gehring, J., Ferraioli, A., Leclére, L., Herl, M., Gao, F., Chevalier, S., Copley, R.R., Houliston, E., et al., 2021. Whole animal multi-plexed single-cell RNA-seq reveals plasticity of *Clytia* medusa cell types. Sci. Adv.7, eabh1683.

Cunningham, K., Anderson, D.J., Weissbourd, B., 2024. Jellyfish for the study of nervous system evolution and function. Curr. Opin. Neurobiol. 88, 102903.

Dereeper, A., Guignon, V. Blanc, G., Audic, S., Buffet, S., Chevenet, F. Dufayard, J.F., Guindon, S., Lefort, V., Lescot, M. Claverie, J.M. Gascuel, O., 2008. Phylogeny.fr: robust phylogenetic analysis for the non-specialist. Nucleic Acids Res. 36, W465–W469.

Emery, M.A., Beavers, K.M., Van Buren, E.W., Batiste, R., Dimos, B., Pellegrino, M.W., Mydlarz, L.D., 2024. Trade-off between photosymbiosis and innate immunity influences cnidarian’s response to pathogenic bacteria. Proc. Biol. Sci. 291, 20240428.

Emery, M.A., Dimos, B.A., Mydlarz, L.D., 2021. Cnidarian pattern recognition receptor repoertoires reflect both phylogeny and life history traits. Front. Immunol. 12, 689463.

Ertűrk-Hasdemir, D., Broemer, M., Leulier, F., Lane, W.S., Paquette, N., Hwang, D., Kim, C.-H., Stőven, S., Meier, P., Silverman, N., 2009. Two roles for the *Drosophila* IKK complex in activation of Relish and the induction of antimicrobial peptide genes. Proc. Natl. Acad. Sci. USA 106, 9779–9784.

Gauthier, M., Degnan, B.M., 2008. The transcription factor NF-κB in the demosponge *Amphimedon queenslandica*: insights on the evolutionary origin of the Rel homology domain. Dev. Genes Evol. 218, 23–32.

Ghosh, G., Van Duyne, G., Ghosh, S., Sigler, P.B., 1995. Structure of the NF-κB p50 homodimer bound to a κB site. Nature 373, 303–310.

Gilmore, T.D., 2006. Introduction to NF-κB: players, pathways, perspectives. Oncogene 25, 6680– 6684.

Gilmore, T.D., Morin, P.J., 1993. The IκB proteins: members of a multifunctional family. Trends Genet. 9, 427–433.

Gilmore, T.D., Wolenski, F.S., 2012. NF-κB: where did it come from and why? Immunol. Rev. 246, 14–35.

Gilmore, T.D., Siggers, T., 2023. NF-kappaB and the immune system. In, *Encyclopedia of Cell Biology, Second Edition* (RA Bradshaw, GW Hart, PD Stahl, eds). Volume 5: 417–426 Oxford, Elsevier

Gold, D.A., Katsuki, T., Li, Y., Yan, X., Regulski, M., Ibberson, D., Holstein, T., Steele, R.E., Jacobs, D.K., Greenspan R.J., 2019. The genome of the jellyfish *Aurelia* and the evolution of animal complexity. Nat. Ecol. Evol. 3, 96–104.

Huxford, T., Huang, D.B., Malek, S., Ghosh, G., 1998. The crystal structure of the IκBα/NF-κB complex reveals mechanisms of NF-κB inactivation. Cell 95, 759–770.

Kane, E.I., Spratt, D.E., 2021. Structural insights into ankyrin repeat-containing proteins and their influence in ubiquitylation. Int. J. Mol. Sci. 22, 609.

Leclère, L., Horin, C., Chevalier, S., Lapébie, P., Dru, P., Peron, S., Jager, M., Condamine, T., Pottin, K., Romano, S., Steger, J., Singaglia, C., Barreau, C., Quiroga Artigas, G., Ruggiero, A. Fourrage, C., Kraus, J.E.M., Poulain, J., Aury, J.-M., Wincker, P., Quéinnec, E. Technau, U., Manuel, M., Momose, T., Houliston, E., Copley, R.R., 2019. The genome of the jellyfish *Clytia hemisphaerica* and the evolution of the cnidarian life-cycle. Nat. Ecol. Evol. 3, 801–810.

Liu H, Zeng L, Yang Y, Guo C, Wang H., 2022. Bcl-3: a double-edged sword in immune cells and inflammation. Front. Immunol. 13, 847699.

Mansfield, K.M., Carter, N.M., Nguyen, L., Cleves, P.A., Alshanbayeva, A., Williams, L.M., Penvose, A.R., Crowder, C., Finnerty, J.R., Weis, V.M., Siggers, T.W., Gilmore, T.D., 2017. Transcription factor NF-κB is modulated by symbiotic status in a sea anemone model of cnidarian bleaching. Sci. Rep. 7, 16025.

Margolis, S.R., Dietzen, P.A., Hayes, B.M., Wilson, S.C., Remick, B.C., Chou, S., Vance, R.E., 2021. The cyclic nucleotide 2’3’-cGMP induces a broad antibacterial and antiviral response in the sea anemone *Nemtatostella vectensis*. Proc. Natl. Acad. Sci. USA 118, e2109022118.

Richter, D.J., Fozouni, P., Eisen, M.B., King, N., 2018. Gene family innovation, conservation and loss on the animal stem lineage. eLife 7. e34226.

Ryzhakov, G., Teixeira, A., Saliba, D., Blazek, K., Muta, T., Ragoussis, J., Udalova, I.A., 2013. Cross-species analysis reveals evolving and conserved features of the Nuclear factor κB (NF-κB) proteins. J. Biol. Chem. 288, 11546–11554.

Schmitz, J.F., Zimmer, F., Bornberg-Bauer, E., 2016. Mechanisms of transcription factor evolution in Metazoa. Nucleic Acids Res. 44, 6287–6297.

Sievers, F., Wilm, A., Dineen, D., Gibson, T.J., Karplus, K., Li, W., Lopez, R., McWilliam, H., Remmert, M., Söding, J., Thompson, J.D., Higgins, D.G., 2011. Fast, scalable generation of high-quality protein multiple sequence alignments using Clustal Omega. Mol. Syst. Biol. 7, 539.

Steele, R.E., David, C.N., Technau, U., 2011. A genomic view of 500 million years of cnidarian evolution Trends Genet. 27, 7–13.

Sullivan, J.C., Finnerty, J.R. 2007. A surprising abundance of human disease genes in a simple “basal” animl, the starlet sea anemone (*Nematostella vectensis*). Genome 50, 689–692.;

Sullivan, J.C., Wolenski, F.S., Reitzel, A.M., French, C.E., Traylor-Knowles, N., Gilmore, T.D., Finnerty, J.R., 2009. Two alleles of NF-κB in the sea anemone *Nematostella vectensis* are widely dispersed in nature and encode proteins with distinct activities. Plos One 4, e7311.

Sun, S.-C., 2011. Non-canonical NF-κB signaling pathway. Cell Res. 21, 71–85.

Williams, L.M., Fuess, L.E., Brennan, J.J., Mansfield, K.M., Salas-Rodriguez, E., Welsh, J., Awtry, J., Banic, S., Chacko, C., Chezian, A., Dowers, D., Estrada, F., Hsieh, Y.-H., Kang, J., Li, W., Malchiodi, Z., Malinowski, J., Matuszak, S., McTigue, T., Mueller, D., Nguyen, B., Nguyen, M., Nguyen, P., Nguyen, S., Njoku, N., Patel, K., Pellegrini, W., Pliakas, T., Qadir, D., Ryan, E., Schiffer, A., Thiel, A., Yunes, S.A., Spilios, K.E., Pinzón C, J.H., Mydlarz, L.D., Gilmore, T.D., 2018. A conserved Toll-like receptor-to-NF-κB signaling pathway in the endangered coral *Orbicella faveolata*. Dev. Comp. Immunol. 79, 128–136.

Williams, L.M., Inge, M.M., Mansfield, K.M., Rasmussen, A., Afghani, J., Agrba, M., Albert, C., Andersson, N., Babaei, M., Babaei, M., Bagdasaryants, A., Bonilla, A., Browne, A., Carpenter, S., Chen, T., Christie, B., Cyr, A., Dam, K., Dulock, N., Erdene, G., Esau, L., Esonwune, S., Hanchate, A., Huang, X., Jennings, T., Kasabwala, A., Kehoe, L., Kobayashi, R., Lee, M., LeVan, A., Liu, Y., Murphy, E., Nambiar, A., Olive, M., Patel, D., Pavesi, F., Petty, C.A., Samofalova, Y., Sanchez, S., Stejskal, C., Tang, Y., Yapo, A., Cleary, J.P., Yunes, S.A., Siggers, T., Gilmore, T.D.,. 2020. Transcription factor NF-κB in a basal metazoan, the sponge, has conserved and unique sequences, activities, and regulation. Dev. Comp. Immunol. 104, 103599.

Williams, L.M., Gilmore, T,D., 2020. Looking down on NF-κB. Mol. Cell. Biol. 40, e00104–20.

Williams, L.M., Sridhar, S., Samaroo, J., Peart, J., Adindu, E.K., Addanki, A., DiRusso, CJ, BB522 Molecular Biology Laboratory, Aguirre Carrión, P.J., Rodriguez Sastre, N., Siggers, T., Gilmore, T.D. 2021. Comparison of NF-κB from the protists Capsaspora owczarzaki and Acanthoeca spectabilis reveals extensive evolutionary diversification of this transcription factor. Commun. Biol. 4, 1404.

Wolenski, F.S., Garbati, M.R., Lubinski, T.J., Traylor-Knowles, N., Dresselhaus, E., Stefanik, D.J., Goucher, H., Finnerty, J.R., Gilmore, T.D., 2011. Characterization of the core elements of the NF-κB signaling pathway of the sea anemone *Nematostella vectensis*. Mol. Cell. Biol. 31, 1076–1087.

Wolenski, F.S., Bradham, C.A., Finnerty, J.R., Gilmore, T.D., 2013. NF-κB is required for cnidocyte development in the sea anemone *Nematostella vectensis*. Dev. Biol. 373, 205– 215.

