## Supplementary Information for "Comparison of Activities of Transcription Factor NF-κB from Two Jellyfish Models"

*bioRxiv*

This file contains Supplementary Tables, Supplementary Figures, and Supplementary References

| Supplemental Table 1. Rel Homology Domain (RHD) Sequences used for phylogenetic analysis (Figure 1B of text). | |
|---|---|
| Organism NF-κB | Amino acid sequences |
| >Homo_sapiens_NF-kB2 (p100) | PYLVIVEQPKQRGFRFRYGCEGPSHGGLPGASSEKG<br>RKTYPTVKICNYEGPAKIEVDLVTHSDPPRAHAHSL<br>VGKQCSELGICAVSVGPKDMTAQFNNLGVLHVTKKN<br>MMGTMIQKLQRQRLRSRPQGLTEAEQRELEQEAKEL<br>KKVMDLSIVRLRFSAFLRASDGSFSLPLKPVISQPI<br>HDSKSPGASNLKISRMDKTAGSVRGGDEVYLLCDKV<br>QKDDIEVRFYEDDENGWQAFGDFSPTDVHKQYAIVF<br>RTPPYHKMKIERPVTVFLQLKRKRGGDVSDSKQFTY<br>YPLVEDKEEVQRKRRK |
| >Homo_sapiens_NF-kB1 (p105) | PYLQILEQPKQRGFRFRYVCEGPSHGGLPGASSEKN<br>KKSYPQVKICNYVGPAKVIVQLVTNGKNIHLHAHSL<br>VGKHCEDGICTVTAGPKDMVVGFANLGILHVTKKKV<br>FETLEARMTEACIRGYNPGLLVHPDLAYLQAEGGGD<br>RQLGDREKELIRQAALQQTKEMDLSVVRLMFTAFLP<br>DSTGSFTRRLEPVVSDAIYDSKAPNASNLKIVRMDR<br>TAGCVTGGEEIYLLCDKVQKDDIQIRFYEEEENGGV<br>WEGFGDFSPTDVHRQFAIVFKTPKYKDINITKPASV<br>FVQLRRKSDLETSEPKPFLYYPEIKDKEEVQRKRQK |
| >Homo_sapiens_RelA | PYVEIIEQPKQRGMRFRYKCEGRSAGSIPGERSTDT<br>TKTHPTIKINGYTGPGTVRISLVTKDPPHRPHPHEL<br>VGKDCRDGFYEAELCPDRCIHSFQNLGIQCVKKRDL<br>EQAISQRIQTNNNPFQVPIEEQRGDYDLNAVRLCFQ<br>VTVRDPSGRPLRLPPVLSHPIFDNRAPNTAELKICR<br>VNRNSGSCLGGDEIFLLCDKVQKEDIEVYFTGPGWE<br>ARGSFSQADVHRQVAIVFRTPPYADPSLQAPVRVSM<br>QLRRPSDRELSEPMEFQYLPDTDDRHRIEEKRKRT |

| | |
|---|---|
| >Homo_sapiens_RelB | PYLVITEQPKQRGMRFRYECEGRSAGSILGESSTEA<br>SKTLPAIELRDCGGLREVEVTACLVWKDWPHRVHPH<br>SLVGKDCTDGICRVRLRPHVSPRHSFNNLGIQCVRK<br>KEIEAAIERKIQLGIDPYNAGSLKNHQEVDMNVVRI<br>CFQASYRDQQGQMRRMDPVLSEPVYDKKSTNTSELR<br>ICRINKESGPCTGGEELYLLCDKVQKEDISVVFSRA<br>SWEGRADFSQADVHRQIAIVFKTPPYEDLEIVEPVT<br>VNVFLQRLTDGVCSEPLPFTYLPRDHDSYGVDKKRK<br>R |
| >Homo_sapiens_cRel | PYIEIIEQPRQRGMRFRYKCEGRSAGSIPGEHSTDN<br>NRTYPSIQIMNYYGKGKVRITLVTKNDPYKPHPHDL<br>VGKDCRDGYYEAEFGQERRPLFFQNLGIRCVKKKEV<br>KEAIITRIKAGINPFNVPEKQLNDIEDCDLNVVRLC<br>FQVFLPDEHGNLTTALPPVVSNPIYDNRAPNTAELR<br>ICRVNKNCGSVRGGDEIFLLCDKVQKDDIEVRFVLN<br>DWEAKGIFSQADVHRQVAIVFKTPPYCKAITEPVTV<br>KMQLRRPSDQEVSESMDFRYLPDEKDTYGNKAKKQ |
| >Mus_musculus_NF-kB2 (p100) | PYLVIVEQPKQRGFRFRYGCEGPSHGGLPGASSEKG<br>RKTYPTVKICNYEGPAKIEVDLVTHSDPPRAHAHSL<br>VGKQCSELGVCAVSVGPKDMTAQFNNLGVLHVTKKN<br>MMEIMIQKLQRQRLRSKPQGLTEAERRELEQEAKEL<br>KKVMDLSIVRLRFSAFLRASDGSFSLPLKPVISQPI<br>HDSKSPGASNLKISRMDKTAGSVRGGDEVYLLCDKV<br>QKDDIEVRFYEDDENGWQAFGDFSPTDVHKQYAIVF<br>RTPPYHKMKIERPVTVFLQLKRKRGGDVSDSKQFTY<br>YPLVEDKEEVQRKRRK |
| >Mus_musculus_NF-kB1 (p105) | PYLQILEQPKQRGFRFRYVCEGPSHGGLPGASSEKN<br>KKSYPQVKICNYVGPAKVIVQLVTNGKNIHLHAHSL<br>VGKHCEDGVCTVTAGPKDMVVGFANLGILHVTKKKV<br>FETLEARMTEACIRGYNPGLLVHSDLAYLQAEGGGD<br>RQLTDREKEIIRQAAVQQTKEMDLSVVRLMFTAFLP<br>DSTGSFTRRLEPVVSDAIYDSKAPNASNLKIVRMDR<br>TAGCVTGGEEIYLLCDKVQKDDIQIRFYEEEENGGV<br>WEGFGDFSPTDVHRQFAIVFKTPKYKDVNITKPASV<br>FVQLRRKSDLETSEPKPFLYYPEIKDKEEVQRKRQ |

| | |
|---|---|
| **C>Mus_musculus_RelA** | **PYVEIIEQPKQRGMRFRYKCEGRSAGSIPGERSTDT<br>TKTHPTIKINGYTGPGTVRISLVTKDPPHRPHPHEL<br>VGKDCRDGYYEADLCPDRSIHSFQNLGIQCVKKRDL<br>EQAISQRIQTNNNPFHVPIEEQRGDYDLNAVRLCFQ<br>VTVRDPAGRPLLLTPVLSHPIFDNRAPNTAELKICR<br>VNRNSGSCLGGDEIFLLCDKVQKEDIEVYFTGPGWE<br>ARGSFSQADVHRQVAIVFRTPPYADPSLQAPVRVSM<br>QLRRPSDRELSEPMEFQYLPDTDDRHRIEEKRKRT** |
| **>Mus_musculus_RelB** | **PYLVITEQPKQRGMRFRYECEGRSAGSILGESSTEA<br>SKTLPAIELRDCGGLREVEVTACLVWKDWPHRVHPH<br>SLVGKDCTDGVCRVRLRPHVSPRHSFNNLGIQCVRK<br>KEIEAAIERKIQLGIDPYNAGSLKNHQEVDMNVVRI<br>CFQASYRDQQGHLHRMDPILSEPVYDKKSTNTSELR<br>ICRINKESGPCTGGEELYLLCDKVQKEDISVVFSTA<br>SWEGRADFSQADVHRQIAIVFKTPPYEDLEISEPVT<br>VNVFLQRLTDGVCSEPLPFTYLPRDHDSYGVDKKRK<br>K** |
| **>Mus_musculus_cRel** | **PYVEIIEQPRQRGMRFRYKCEGRSAGSIPGERSTDN<br>NRTYPSVQIMNYYGKGKIRITLVTKNDPYKPHPHDL<br>VGKDCRDGYYEAEFGPERRPLFFQNLGIRCVKKKEV<br>KGAIILRISAGINPFNVGEQQLLDIEDCDLNVVRCV<br>FMFFLPDEDGNFTTALPPIVSNPIYDNRAPNTAELR<br>ICRVNKNCGSVRGGDEIFLLCDKVQKDDIEVRFVLN<br>DWEARGVFSQADVHRQVAIVFKTPPYCKAILEPVTV<br>KMQLRRPSDQEVSESMDFRYLPDEKDAYGNKSKKQK** |
| **>Gallus_gallus_NF-kB2 (p100)** | **PYLVIIEQPKQRGFRFRYGCEGPSHGGLPGASSEKG<br>HKTYPTVKICNYEGMARIEVDLVTHSDPPRVHAHSL<br>VGKQCNEAGNCVAIVGPKDMTAQFSNLGVLHVTKKN<br>MMEIMKEKLKKQKTRNTNGLLTEAELREIELEAKEL<br>KKVMDLSIVRLRFTAYLRDSSGNFTLALQPVISDPI<br>HDSKSPGASNLKISRMDKTAGSVRGGDEVYLLCDKV<br>QKDDIEVRFYEDDENGWQAFGDFSPTDVHKQYAIVF<br>RTPPYHKPKIDRPVTVFLQLKRKKG** |

| | |
|---|---|
| >Gallus_gallus_NF-kB1 (p105) | PYLQIIEQPKQRGFRFRYVCEGPSHGGLPGASSEKN<br>KKSYPQVKICNYVGPAKVIVQLVTNGKYVHLHAHSL<br>VGKFCEDGVCTVNAGPKDMVVGFANLGILHVTKKKV<br>FETLETRMIDACKKGYNPGLLVHPELGYLQAEGCGD<br>RQLTEREREIIRQAAVQQTKEMDLSVVRLMFTAFLP<br>DSNGGFTRRLDPVISDAIYDSKAPNASNLKIVRMDR<br>TAGCVTGGEEIYLLCDKVQKDDIQIRFYEEDENGGM<br>WEGFGDFSPTDVHRQFAIVFKTPKYRDVNITKPASV<br>FVQLRRKSDLETSEPKPFLYYPEIKDKEEVQRKRQK<br>L |
| >Gallus_gallus_RelA | PFVEILEQPKQRGMRFRYKCEGRSAGSIPGEHSTDS<br>ARTHPTIRVNHYRGPGRVRVSLVTKDPPHGPHPHEL<br>VGRHCQHGYYEAELSPERCVHSFQNLGIQCVKKREL<br>EAAVAERIRTNNNPFNVPMEERGAEYDLSAVRLCFQ<br>VWVNGPGGLCPLPPVLSQPIYDNRAPSTAELRILPG<br>DRNSGSCQGGDEIFLLCDKVQKEDIEVRFWAEGWEA<br>KGSFAAADVHRQVAIVFRTPPFRERSLRHPVTVRME<br>LQRPSDRQRSPPLDFRYLPHQGDLQCIEEKRKRT |
| >Gallus_gallus_RelB | PRLIITEQPKKTGMRFRYECEGRSAGSILGESSTEA<br>SKTLPAIELLNCQAIPEVQVTACLVWKDWPHRVHPH<br>GLVGKDCSNGLCQVRLQPHANPRHSFSNLGIQCVKK<br>KEIEAAIEKKLQLGIDPFKAGSLKNHQEVDMNVVRI<br>CFQASYRDGSGRTRQLSPVLSEPIFDKKSTNTSELR<br>ICRMNKESGPCTGGEELYLLCDKVQKEDIAVVFRKE<br>PWEARADFSQADVHRQGAIVLRTPPYRCVQLSEPVQ<br>VEVFLQRLTDRARSRGCPYTYLPRERDAYGVKVKRK<br>RGMPDLLEELSGADPYGIEAKRRKP |
| >Gallus_gallus_cRel | PYIEIFEQPRQRGMRFRYKCEGRSAGSIPGEHSTDN<br>NKTFPSIQILNYFGKVKIRTTLVTKNEPYKPHPHDL<br>VGKDCRDGYYEAEFGPERRVLSFQNLGIQCVKKKDL<br>KESISLRISKKINPFNVPEEQLHNIDEYDLNVVRLC<br>FQAFLPDEHGNYTLALPPLISNPIYDNRAPNTAELR<br>ICRVNKNCGSVKGGDEIFILCDKVQKDDIEVRFVLD<br>NWEAKGSFSQADVHRQVAIVFRTPPFLRDITEPITV<br>KMQLRRPSDQEVSEPMDFRYLPDEKDPYGNKAKRQR |

| | |
|---|---|
| >Capsaspora_owczarzaki | DLLMVTEEPAQFARFRYMSEQRERSLAGENSFPTLM<br>VNPKYARVVPEMALVTAVLVTKMPDPHTGRQQKHWH<br>HLGGIPAAPLEGPQRIARFDNIAVIMDKANNKDKDK<br>SKAPVRSKDDQRCVRIMFELVFVSGNTQFYGRAISQ<br>PIYNAKLAITKISHSSGPVTGGNEVIMLCSKIRKGV<br>TGVRMTDPTQWSVQAPSGSAWELNPQTLKADCNVPG<br>ANLFFHHQYAVVLTLPPYHTQTITAPVTVRISILDT<br>DDETESQYVEYTYLPAEAAVRNAELAARKRRR |
| >Amphimedon_queenslandica | LEIVEQPKSRGFRFRYDCEGQSHGGLPGENSEKNRR<br>QKTYPTVHLKGYRGRARVMVSLVTDSDPAMPHAHSI<br>VGKNAIDGRCVVEIGPETDMYAQFTSLGILHVTKKK<br>VPEVLTRRLLQQTTPRGQMVDQMEVVDVDMTTAQLT<br>SEEQDEIHQQAQTLAKSMNLSVVRLCFQAFLPDENG<br>RYTIPIDPVFSNKVYDSKAPSAGTLKICRLDRTSGS<br>VKGGDDVFLLCDKVQKNDIEVVFYEDKQETTGGMQL<br>QPWMAKGRFGPNDVHHQYAIVFQTPTFYNQAIEHPV<br>QVWIALKRPSDHETSEPKPFLYLPQEFDEERIGQKR<br>RKK |
| >Nematostella_vectensis | PYLEILEQPKPRGFRFRYPSEGPSHGGLPGQFSTSK<br>SKSYPSVQVNNYQGPCRIVVTLVTKDEPYMLHAHSL<br>TGKNANEEGVVTVQVGPDQHMTASFPNLGIQHVTKK<br>NVVKVLMDRFIKWQTLQNATFAKLSEGIKDGVDLSL<br>FGVNTAINSNKLGFDKNVALSVANQEAAKSREYAKQ<br>QAAAMDLSAVRLCFQAYLPDQDGNFTRPLKPVYSDA<br>VLDSKEPSASQLKICRMDKNSGCVTGGDEIYLLCDK<br>VQKDDIEIHFYEMDDITGKYTWEDLGKFSPCDVHRQ<br>FAIVFKTPPYWNIAIERPANVLVELRRKKK |
| >Exaiptasia_pallida | PYLEILEQPKSRGFRFRYPCEGPSHGGLPGEFSDSK<br>NKSYPSVQVCNYQGPCRIVVSLVTEDEPHMPHAHSL<br>TGKHANNDGIVTVQIGTEQGMTASFPNLGIQHVTKK<br>KVAKTLTERYTKMQALQNATLTALATNNSTPSSFMN<br>FGSVAREQVMASQGPFDRNLAAAVAGEETKKILKLV<br>QEQSKTMNLSAVRLCFQAYLPDENGNFTKPLKPCIS<br>NPVYDSKAPASCQLKICRMDKNSGCVTGGDEIYLLC<br>DRVQKDDIEIRFYENNDDGKPIWEDTGKFAPADVHR<br>QFAIVFKTPAYHNIAIERPVEVLLELRRKSDKETSE<br>PFTFTYSPQMFDTEQIGAKRRK |

| | |
|---|---|
| >Orbicella_faveolata | PYMEILEQPKQRGFRFRYPCEGPSHGGLPGQYSEKG<br>KKSYPSVQLCNYHGPARIVVSLVTVDEPPMPHAHSL<br>IGKNSNNGAVTVQIGPEHGMTASFPNLGIQHVTKKS<br>VGKVLMERYIKMQTLHTATLHALTADSKGFDMELVG<br>DQALADGETATFNRTMAEAVAAEESQKVRQMVEDQK<br>QSMNLNAVRLCFQAYLPDDGGCFTKALPPCISNPVY<br>DSKAPSASNLKICRMDRNSGCVTGGDEVYLLCDKVQ<br>KEDIDVMFYEIDVETGKKTWEAGGVFAPTDVHRQVA<br>IVFKTPAYWNIATERPVKVHLELRRKSDQETSEPVE<br>FTYQPQLFDKEQIGAKRRKK |
| >Aurelia_aurita | PYLEILLQPKQRGYRFRYNSEGITHGGIPAESTEKG<br>NKKFPTVHIANYKGKAAICITLVTAEDPPQVHAHNL<br>VGKDVTKGMLYCEDDKQEWVKSFTNLGIQHVTKKDL<br>VKVLHEKLCQSYQLTQFSAAIKTEDETQSFDISALI<br>GSFGGTDGAGTMIDESVAMAVAEEENKRLRKEAEEM<br>AKTIDLSAVRLCIVAYLPDEAGLLTRALPAVFSKPI<br>FDQKAAHSGQLKICRISKPSSSVNGGEEVFLLCDKV<br>QKDDISVRFYQEDGAGNATWEGFGKFTQADVHKQSA<br>IVFTTPPYVDKAIQRPVEVWLQLKRGKDKECSEPVK<br>FTYRPEEYDRYKIGEKRRKG |
| > Clytia_hemisphaerica | PYPFKGAARLEITRQPKKRGYRFRYLSEGKTHGMLP<br>GEPSDSGEKVFPSVRIVNHLGNAKVVMYLVTEEDPQ<br>FIHPHTLLIDKIPVGGYHIFDVNEDFDVQLKNVAIQ<br>HVNKQDLPTKMLDRHLQSKYIKEVYSMNQYGANPKI<br>DIDTFASNLADKAKKQPFFDHRTKQALTGEEEIQLS<br>LAMRKKQKEINMSSCRLCFIAFLQDPETGKFDKILQ<br>PVHSDLIIDGKKKEGAPLKIIRVSHVAGSVEGGKEV<br>WLLSDKIDAEDTEVYFWERSQDKQEMFWEGFGEFNK<br>TDVYKQALIVFKIPPYCNQNIDQPRTVNLQLRRKKD<br>RNCVSDVHHFSYKPKHYDRYGLREKRRKN |
| >Cassiopea_xamachana | PYLEILRQPKQRGYRFRYSSEGNTHGGIPAVCSEKG<br>QKTFPTVQIANYSGKAAICVSLVTVDDPPRVHAHNL<br>VGKDTSNGLFYIEDTKKSWTHSFQNLGIQHVTKKDL<br>ITVLHNKLCQQYQLTLIPQTKPDEAFQSFDIAALVD<br>SFGAVDGPSTIFDEPMAKLVENEKKQLYAEAEKLAK<br>SIDLSAVRLCFQAFLPDESEKLTRSLPPVYSVPIYD<br>QKAAHAGQLRICRISSSSSSVNGDEEVFLLCDKVQK<br>DDIAVQFFTEEGKITWEGFGKFSHSDVHKQCAIVFR<br>TPPYIDKAIQRPVDVFVRLRRGKDNEFSPAIPFTYK<br>PETFDKYKIGEKRRKG |

| | |
|---|---|
| >Hydra_vulgaris | PYLKIERQPRKYGYRFRYKTEGVCHGGILADTDGAV<br>NCGSSKSCPKIKVHNLNGQRAKVVLRLAAEHDNETM<br>HIHSLVYNKTVTNGVHLLTLNEVDEVELEHVAVQQE<br>KTKETKNFLFKERVLQSEYLKKYNSINPPIDVETFV<br>KRLEDREDKKLINRLTGVECNNKDLQSEASSLMKDL<br>NVHSVVLAFHCFLEDSNGRYTINLPTVYSVPIYDGK<br>NKKGCEYKILRLSSVSGTPKGGEEVWMLCDKFDKSD<br>VEVRFFDDTPVVPWSALAIVNKSDVHNQNLIIFRTP<br>PYKHKVLDQPVKVKIELRAASDHMRCSKHFDYTYNN<br>SSDDGYEVGKKRKK |
| >Rhopilema_esculentum | PYIEILKQPKQRGYRFRYISEGQTHGGITADNTNKGNKQY<br>PSIHIGNYKGRASVCITLVTADDPVRVHPYNLVGKDVKNG<br>MFHCDETKEQWVRSFPNLGIQNVTKKDLVRVMHEKLCQQF<br>QLNQLSAAKDNSEITQSFDIAHLLESFGGNDEPGLAIDES<br>VALVIAEEEKRRLLIEAEKISKEIDLSAVRLCFQAFLLDE<br>SGQITKTLPAVLSQPIYDQKAAHAGQLRICRISKTSSSVA<br>GGEEVFLLCDKVQKEHIAVRLYQEDSAGKLTWEGFGKFSQ<br>ADVHKQSAIVFRTPPYVDKAIQKPIEVKLQLKRSKDNACS<br>EPVPFIYQPVEYDQYKIGEKRRKG |
| >Hydractinia_symbiolongicarpus | PFLVIERQPKKRGYRFRYSSEGQTHGGIPGEPDKSN<br>DNRKVYPSLRLHGHNGQAKVVVTLVTAEDNQPHLHA<br>HSLIVNKVMTKGFHVYEMNGESSLELKSIAIQHVVK<br>ENLLSTLMDRILQTRFVETFGWNSGLNVNGTREEMF<br>DVSHYVQSLMDRHDDTVFYDRATAQAIAGVELENLR<br>KKANDLAKKMQMNQCRLCFQAFLPDSHGKFTRILNP<br>VYSDIIYDGKTKQGAMLKIIRLSKVSCSVKGDEEVW<br>LLADKLNPDDIEVRFFEKETKLSTSMVPWEDTGKFN<br>KTDIYKQVCLIFKTPMYHDQNIRQPANVFIQLRRKT<br>DTSCCSEPMEFKYTPIQYDRYGIGEKRKKE |
| >Xenia_Sp | AELEILEQPKQRGFRFRYSCEGPSHGGLPGENSVKGKKTS<br>PTVQIVNYHGKARVEVSLVTTSDPPTPHAHSLVGKNVDGA<br>CVVEVSEDTGMTAVFQNLGIQHVTKKNVKNVLKERHLKRL<br>SLEKTSISQSSNLLLNVHTNVDSERIAEVITNEEIAKIDR<br>IVEQEAKNMNLSSVRLCFQAYLPNENGYFETCLNPVYSNP<br>IYDSKAAAAAELKICRVDKQSGFVQGGEEVFLLCDKVQKE<br>DIEVRFYEEHTDDDIANIKEPWEALAKFSHTDVHRQFAIV<br>FKTPEYWNTAIEKPVTVLMELRRKSDNERSAPFEFVYKPR<br>EFDTDQIGAKRRKT |

| | |
|---|---|
| > Pocillopora_damincornis | PYMEILEQPKQRGFRFRYPCEGPSHGGLPGQYSEKGKKSY<br>PSVQLCNYQGPARIVVSLVTVDEPPMPHAHSLISKNSNNG<br>VVTVQIGPEQGMTATFPNLGIEHVTKKMVSKVLMDRYIKM<br>QTLHTATLNALTSGDGKVFGVAGLVDQAMVDGDRGSFDKR<br>LAEAVAEEESQKVRAMVEEQKQSMNLNAVRLCFQAFLPDE<br>TGAFTKALPPCISNAVYDSKAPSASNLKICRMDRNSGCVK<br>GGDEVYLLCDKVQKDDIEVIFYETEMDTGKKTWEDRGVFS<br>PTDVHRQVAIVFKTPPYWNVAIEQPVKVQLELRRKSDQET<br>SDPVEFTYQPQMFDNEQIGAKRRKI |
| >Montipora_capricornis | PYLEILEEPKQRGFRFRYPCEGPSHGGLPGQFSEKGKKSY<br>PSVQLCNYQGPARIVVSLVTSDEPPMPHAHSLIGKNSNNG<br>VVIVQIGPEHGMTASFPNLGIQHVTKKKVSQVLMERYLKM<br>QTLHTATLNAMTADNSHFDVGALGDHATSDGDRSIFDKNL<br>AEAIAEEEAKKVRRLVEEQENSMNLSVVKLCFQAFLPDDS<br>GCFTKALPPCFSLPVYDSKAPSAANLKICRMDRNSGSVTG<br>NDEVYLLCDKVQKDDIEVVFYELDPDNGKRTWENRGLFAP<br>TDVHRQVAIVFKTPAYWNVCIERPVKVHLELRRKSDKETS<br>ESVDFTYQPQEFDKEQIGAKRRKK |
| >Acropora_cervicornis | PYLEILEQPKQRGFRFRYPCEGPSHGGLPGEYSEKGKKSY<br>PSVQLCNYKGPARIVVSLVTCDENPMPHAHSLIGKNASNG<br>VVTVQIGPEQGMTASFPNLGIQHVTKKNVGYVLMDRYLKM<br>QMLHTATLNALTTDPRVFDIGAMVDEATADGDRGEFDKQI<br>ADTIAEEEASKVRSLVEKQKNSMNLSVVRLCFQAYLPDDN<br>GCFTKALPPCFSRSVYDSKAPSAANLKICRMDRNSGCVTG<br>SDEVYLLCDKVQKDDIAVVFYEIDENGKRSWEGLGLFAPT<br>DVHRQVAIVFKTPAYWNIAVERPVKVHLELRRKSDDETSD<br>PVEFTYQPQMFDKEQIGSKRRKK |
| >Corticium_candelabrum | PALDIFEQPKSRGFRFRYPCEGPSHGGLPGASSTKSRKSF<br>PTVQLFNYEGRARIVVSLVTLDNPPRPHAHSLVGKNTRNG<br>QCIVELGPEQKMSASFPNLGVLHETKKNVPRALLFRYAWD<br>KFSQLERAGSLPQDLIQQAAAAVPEIDGEVVDVENRPLPP<br>LPEEVMQPLRKQADEDAKNMNLSVVRLCFQAFLPDDSGSF<br>TRALQPCLSDPVYDSKAPSASTLKICRLDRNSGGVNGGDE<br>VFMLCDRVQKDDIDVKFFDKNSSLSQSASLWEANASFSPN<br>DVHRQYAIVFKTPRYWNISITRPVEVFMQLRRRSDGEVSE<br>PKVFTYQPAEYDAEQIGRKRKKR |

| | |
|---|---|
| **>Sycon_ciliatum** | **AELTILEEPRQRGFRFRYSCEGPSHGGLPGETSEKNRKSY<br>PTVQLRKYEGRAARVVVSLVTQDDPPRPHAHSLVGRHVVN<br>GQCTVQIGPETNWMASFPNLGILHVTKKNVVKVLLERYAL<br>ALTGPGASSVALPIMPDLKPGTAPTSSLPDEAAVIAASME<br>KLSEEERSQLKAMAEEASRHMNLSVVRLCFQAYLADEQGS<br>FTKPLQPCISQPVYDSKAPSASTLKISRMDRHSGSVLGGD<br>EVYLLCDRVQKDDIEVRFFTEGDGSDSGDGRPDGWSALGV<br>FAPSDVHRQFAIVFKTPAYCDTSISRPVNVWIQLRRRSDN<br>ETSEAKPFTYQPQIVDTEQLARKRQKK** |
| **>Acropora_muricata** | **PYLEILEQPKQRGFRFRYPCEGPSHGGLPGEYSEKGKKSY<br>PSVQLCNYKGPARIVVSLVTSDENPMPHAHSLIGKNASNG<br>VVTVQIGPEQGMTASFPNLGIQHVTKKNVGLVLMDRYL<br>KMQMLHTATLNALTIDPRVFDIGAMVDEATADGDRGEFDK<br>QIADTIAEEEASKVRSLVEKQKNSMNLSVVRLCFQAYLPD<br>DNGCFTKALPPCFSRSVYDSKAPSAANLKICRMDRNSGCV<br>TGNDEVYLLCDKVQKDDIAVVFYEIDENGKRSWEGQGLFA<br>PTDVHRQVAIVFKTPAYWNIAVERPVKVHLELRRKSDDET<br>SDPVEFTYQPQMFDKEQIGSKRRKK** |
| **>Pocillopora_verrucosa** | **PYMEILEQPKQRGFRFRYPCEGPSHGGLPGQYSEKGKKSY<br>PSVQLCNYQGPARIVVSLVTVDEPPMPHAHSLIGKNSTNG<br>VVTVQIGPEQGMTATFPNLGIEHVTKKMVGKVLMDRYIKM<br>QTLHTATLNALTSGDGKVFGVAGLVDQAMVDGDRSSFDKR<br>LAEAVAEEESQKVRAMVEEQKQSMNLNAVRLCFQAFLPDE<br>TGAFTKALHPCISNAVYDSKAPSASNLKICRMDRNSGCVK<br>GGDEVYLLCDKVQKDDIEVIFYETEMETGKKTWEDRGVFS<br>PTDVHRQVAIVFKTPAYWNVAIEQPVKVQLELRRKSDQET<br>SDPVEFTYQPQMFDKEQIGAKRRKK** |
| **>Montipora_foliosa** | **PYLEILEEPKQRGFRFRYPCEGPSHGGLPGQFSEKGKKSY<br>PSVQLCNYQGPARIVVSLVTSDEPPMPHAHSLIGKNSNNG<br>VVIVQIGPEHGMTASFPNLGIQHVTKKKVSQVLMERYLKM<br>QTLHTATLNAMTADNSHFDVGALGDHATSDGDRSIFDKNL<br>AEAIAEEEAKKVRRLVEEQENSMNLSVVKLCFQAFLPDDS<br>GCFTKALPPCFSLPVYDSKAPSAANLKICRMDRNSGSVTG<br>NDEVYLLCDKVQKDDIEVVFYELDPDNGKRTWENRGLFAP<br>TDVHRQVAIVFKTPAYWNVCIERPIKVHLELRRKSDKETS<br>ESVDFTYQPQEFDKEQIGAKRRKK** |

| | |
|---|---|
| **>Stylophora_pistillata** | **PYMEILEQPKQRGFRFRYPCEGPSHGGLPGQYSEKGKKSY<br>PSVQLCNYQGPARIVVSLVTVDEPPMPHAHSLIGKNSAGG<br>VVTVQIGPEQGMTATFPNLGIEHVTKKMVGKVLMDRYIKM<br>QTLHTATLNALTSADGKGFDVTVLGDHVDGDRSSFDKHLA<br>EAVAEEESQKVRTMVEEQKQSMNLNAVRLCFQAFLPDESG<br>AFTKALHPCISNPVYDSKAPSASNLKICRMDRNSGCVSGS<br>DEVYLLCDKVQKDDIEVIFYETEMETGKKTWEDRGVFAPT<br>DVHRQVAIVFKTPAYWNVAIEQPVKVQLELRRKSDQETSD<br>PVEFTYQPQLFDKEQIGAKRRKK** |
| **>Dendronephthya_gigantea** | **AELEILEQPKQRGFRFRYSCEGPSHGGLPGERSAKGRKSF<br>PSVQIINYSGKARIEVTLVTVSDPPQPHAHSLVGKNVRDG<br>ACIVEVGPDCGMTASFPNLGVQHVTKKNVLPILKERYLKK<br>HRLEKSFNSGVSSGDQVMHVDFHDPASSGAIAEAITNEEM<br>KSIEVIAKQEADKMNLSTVRLCFQTYLPDQDGRFVICLKP<br>VYSHPIYDSKAAAASELKICRIDRQSGFVQGGEEIFLLCD<br>KVQKEDIEVRFFENQEEDRPEPWEAFGKFSTSDVHRQFAI<br>VLKTPEYWNAAIEKPVKVLLELRRKSDKETSAPVEFTYKP<br>QEFDPEQIGAKRRKK** |

**Supplemental Table 2. Plasmids used in this study**

Mainly expression vectors for use in tissue culture and yeast cells

| Plasmid Name | Plasmid Description |
|---|---|
| pUC57-Ch-NF-κB | pUC57-Simple with Ch-NF-κB cDNA codon-optimized for expression in human cells. Synthesized by GenScript. |
| pUC57-Aa-NF-κB | pUC57-Simple with Aa-NF-κB cDNA codon-optimized for expression in human cells. Synthesized by GenScript. |
| pUC57-Ch-IκB | pUC57-Simple with Ch-IκB cDNA codon-optimized for expression in human cells. Synthesized by GenScript. |
| pUC57-Ch-BCL-3 | pUC57-Simple with Ch-BCL3 cDNA codon-optimized for expression in human cells. Synthesized by GenScript. |
| pUC57-Aa-BCL3 | pUC57-Simple with Aa-IκB cDNA codon-optimized for expression in human cells. Synthesized by GenScript. |
| pUC57-Aa-NF-κB | pUC57-Simple with Aa-NF-κB cDNA codon-optimized for expression in human cells. Synthesized by GenScript. |
| pUC57-Ch-IκB | pUC57-Simple with Ch-IκB cDNA codon-optimized for expression in human cells. Synthesized by GenScript. |
| pUC57-Ch-BCL-3 | pUC57-Simple with Ch-BCL3 cDNA codon-optimized for expression in human cells. Synthesized by GenScript. |
| pUC57-Aa-BCL3 | pUC57-Simple with Aa-IκB cDNA codon-optimized for expression in human cells. Synthesized by GenScript. |
| GBT9 | Ref. 1 |
| GB-Nv-NF-κB | Ref. 1 |

| | |
|---|---|
| GB-Ch-NF-κB | A linker-based BamHI-BamHI fragment encoding aa 2-477 of Co-NF-κB was subcloned into BamHI-digested pGBT9 vector. |
| GB-Aa-NF-κB | A linker-based BamHI-BamHI fragment encoding aa 2-435 of Aa-NF-κB was subcloned into BamHI-digested pGBT9 vector |
| GB-Ch-BCL-3 | A linker-based BamHI-BamHI fragment encoding aa 2-739 of Ch-BCL3 was subcloned into BamHI-digested pGBT9 vector. |
| pcDNA MYC vector | Ref. 1 |
| MYC-Ch-IκB | EcoRI-XbaI fragment encoding amino acids 2-297 of Aa-IκB was subcloned into EcoRI-XbaI digested pcDNA-MYC vector. |
| MYC-Ch-BCL3 | EcoRI-XbaI fragment encoding amino acids 2-739 of Ch-BCL3 was subcloned into EcoRI-XbaI digested pcDNA-MYC vector. |
| MYC-Aa-BCL3 | EcoRI-XbaI fragment encoding amino acids 2-651 of Aa-BCL3 was subcloned into EcoRI-XbaI digested pcDNA-MYC vector. |
| 3X-κB-luc | Ref 1 |

**Supplemental Table 3. Innate immune pathway homologs in *A. aurita* and *C._hemisphaerica***

| **Protein Name** | **Homolog Present in *C. hemisphaerica*** | **Accession Number of *C. hemisphaerica* Homolog** | **Homolog Present in *A. aurita*** | **Accession Number of *A. aurita* Homolog** |
|---|---|---|---|---|
| TLR | + | XP_066922741.1, TCONS_0005672<br><br>TCONS_00000159 | + | scaffold128.g8 |
| IL-1R | - | - | + | scaffold8.g29, scaffold1880.g2 |
| MYD88 | -* | - | +* | Seg116.2 |
| IRAK | -* | - | -* | - |
| TRAF | +* | XP_066932459.1<br>XP_066918860.1<br>XP_066926495.1<br>XP_066925993.1<br>XP_066918082.1<br>XP_066935920.1 | +* | scaffold3.g45, scaffold246.g18, scaffold347.g10, scaffold1411.g5, scaffold308.g16, scaffold308.g17, scaffold3474.g3 |
| NEMO/IKKγ | +* | XP_066910580.1 | +* | scaffold301.g3 |
| IKKα | -* | - | +* | scaffold634.g1 |
| IKKβ | +* | XP_066933320.1 | -* | - |
| TBK/IKKε | +* | TCONS_00026766 | - | - |
| IκB | +* | XP_066933946.1<br><br>XP_066933186.1, TCONS_00042973-protein | - | - |
| NF-κB | +* | XP_066928423.1, TCONS_00005923 | + | scaffold238.g7 |
| BCL-3 | + | TCONS_00046043 | + | scaffold146.g27 |
| RAC1 | +* | TCONS_00065695 | -* | - |
| PI3K | +* | TCONS_00002349, TCONS_00002351 | +* | scaffold935.g2, scaffold2153.g5, scaffold863.g3, scaffold790.g12 |

| | | | | |
|---|---|---|---|---|
| AKT | +* | TCONS_00027365 | -* | - |
| TRIF | - | - | - | - |
| TRAM | -* | - | +* | scaffold162.g6 |
| RIPK | -* | - | - | - |
| TOLLIP | - | - | + | scaffold114.g6 |
| Ubc13 | + | TCONS_00003004 | + | scaffold1623.g6 |
| ECSIT | - | - | - | - |
| UEV1A | - | - | - | - |
| TAK1 | + | TCONS_00027570, TCONS_00027576 | + | scaffold583.g13, scaffold16.g5 |
| TAB 1/2 | +* | TCONS_00058564 | -* | - |
| A20 | - | - | - | - |
| CTLs | +* | TCONS_0000809, TCONS_00009141 | +* | scaffold468.g11 |
| SYK | +* | TCONS_00049928 | +* | scaffold383.g12 |
| CARD9 | -* | - | -* | - |
| BCL10 | -* | - | -* | - |
| MALT1 | +* | TCONS_00022168, TCONS_00036912, TCONS_00073355 | +* | scaffold238.g3 |
| CASP8 | +* | TCONS_00063807 | -* | - |
| CASP10 | +* | TCONS_00018475, TCONS_00061067 | -* | - |
| RIG1 | - | - | + | scaffold111.g19 |
| MDA5 | - | - | + | scaffold494.g1 |
| MAVs | - | - | -* | - |
| TRADD | -* | - | -* | - |
| FADD | -* | - | -* | - |
| RIP1 | - | - | -* | - |
| MEKK1 | +* | TCONS_00013759 | +* | scaffold130.g23 |
| NLRs | +* | TCONS_00001838, TCONS_00031619, TCONS_00058987, | -* | - |

| | | | | |
|---|---|---|---|---|
| | | TCONS_00059382, TCONS_00072663 | | |
| RIP2 | -* | - | -* | - |
| TIRAP/MAL | -* | - | -* | - |

*Indicates a homolog identified by Emery et al. (2021)

**Supplemental Figure 1**

# A

**Ch-NF-κB (477 aa)**
MNLPEVTKNPIYGSSMIGASYPMSMSVPYPFKGAARLEITRQPKKRGYRFRYLSEGKTHG
MLPGEPSDSGEKVFPSVRIVNHLGNAKVVMYLVTEEDPQFIHPHTLLIDKIPVGGYHIFD
VNEDFDVQLKNVAIQHVNKQDLPTKMLDRHLQSKYIKEVYSMNQYGANPKIDIDTFASNL
ADKAKKQPFFDHRTKQALTGEEEIQLSLAMRKKQKEINMSSCRLCFIAFLQDPETGKFDK
ILQPVHSDLIIDGKKKEGAPLKIIRVSHVAGSVEGGKEVWLLSDKIDAEDTEVYFWERSQ
DKQEMFWEGFGEFNKTDVYKQALIVFKIPPYCNQNIDQPRTVNLQLRRKKDRNCVSDVHH
FSYKPKHYDRYGLREKRRKNLPPEIYESTPPIKRQHMDYAADSPPASTASKYQPYDSAHI
NNPTLSDIVMPQGRNEYRAIVDASTRHSPAVPTTFNGPLDNYRKNHHSSKESDTDEN

**Aa-NF-κB (435 aa)**
MNIGNDIDVPEELLQQILTGNYDFNMKTDDEPYLEILLQPKQRGYRFRYNSEGITHGGIP
AESTEKGNKKFPTVHIANYKGKAAICITLVTAEDPPQVHAHNLVGKDVTKGMLYCEDDKQ
EWVKSFTNLGIQHVTKKDLVKVLHEKLCQSYQLTQFSAAIKTEDETQSFDISALIGSFGG
TDGAGTMIDESVAMAVAEEENKRLRKEAEEMAKTIDLSAVRLCIVAYLPDEAGLLTRALP
AVFSKPIFDQKAAHSGQLKICRISKPSSSVNGGEEVFLLCDKVQKDDISVRFYQEDGAGN
ATWEGFGKFTQADVHKQSAIVFTTPPYVDKAIQRPVEVWLQLKRGKDKECSEPVKFTYRP
EEYDRYKIGEKRRKGLPENLEEMLNTAKQSRPATSTISSAPADLSQFNMPEGGPLDISGI
RFFKQSNQGPFSSRK

**Ch- IκB (293 aa)**

MSNPKDKLPNQPIDSGIGDSFGPDFVPDDEKRLTEELENLNIQPSNQINTKTLLDQAFTPD

EEND**TYLH**VYIAKNSPDEAMQLIEMCPDKNLLNIQNYLGQ**TPLH**VASYVNLDKVAMSLVHH

DANLEIQDRDGKNVFHICAERGHIETLENVIKMACQTNKTNSAWNLLHSTDFEGQAPFFLA

ALNKKKQMCVTLAKLNIDVNQIDIKNGN**TPIH**EAILEQNIDYDFLEFLVKTCNLNINAQNY

AGI**TALH**LAAGRNDSNTYSKLLELGANDSFVDIRDFTPEMCGPSDSIQLLNGV

**Ch Bcl-3 (727)**

MPKPYSNHSKKAEENTERFQKALQSKMAQSVATLKTTKVEGLSNLVDAARFVEKTNSSKIT

KKAKPKPIYKGKERGKEKKSHRESKHLIVEDTDADDETSDIESEVESDSQLALLKKSVSEM

MAKIQRLESKRKRKKMPKKAHLSDYEDSSDEDVKPKKPLKKESSPPTKPINANQQLTMHSQ

QMMQMQQQYWQMQQMMNMYSNPSINPAGMMPSMPFMTTQHQQQPSFVNMNSKSASNTLRFP

KTKKPNAETRFKDSSKIEDKKSRKSVEGEAGEETDASEVYAPTAIKDPINEYLGKDPFVPT

PSFISFVDDITKDEKTVQSTVVSSTKRAAETSKVPGSDIPRALFQRNDSITNTQQISATNQ
GNPSYAGILMNLFPTGKPNPLEIRENDKRSEEINTVLRSMIGYNPVDAKPKQPKRFDLSAD
RHTHSKPLFQNLLPISKHSQSPIPSDNAPESKRFVDNATWLRECSKKMDLQDQEGD**TILH**I
LIAKEETPKAIEVIKKMQHLPSLDILNALGQ**TPLH**LAMYTKNLPVVKELLQHGSDVSLVDA
HGN**NPLH**IACEENSIEMLEIVFNEGLSHYQTATNTSAMITTLAPEYFHIINARNNQGL**AAL**
**H**IAAKEDNEVIIKFLGERGAEMNNQEGRSGN**TPLL**IALMKNNWKMASFLLERFVNVNIPNF
SNF**YPLH**FAVQGNHVEMVKALLSRGAEMGSRANDYDQNANTTEIKQLLAKEMRRRRRNRIK
LEPKSQM

**Aa-BCL3 (641 aa)**

MGDGREGQSLLQRDKEGDTFLHIATARNEHSYIEKILRTDVAKELINAQNNHGQ**TALH**IAT
IMGQGDLVKLLLQYGAILQLQNNDGD**TSIH**LATKLGKIVCLMILCENVTKEILNIGNDSGE
**VPLH**LAAKSENLQAFGLFLQKEADVKFRNEKTGM**TALH**FAVEKENKEMVEKLLELGADVNA
QATNGNTPLHSLLGAEQRNTVKLLLDKGADINAKNEAGQSPHDLADKTMQRFMTRNRKDRK
TRRPKRPKGATASDNQPEPETSTTPKPVSDPTHSNEAQLSKRPFIPLTRVQSHPGKKPRLF
EDGGASNTSQGSAAPAQRQLTSSASVDHCVQLDAWGRRLDSQSLSRGNSDEPCDDIERQEC
LKLPFTAYKSMSMPEFSNGKVDVTNSIFGQQQQTLTRFQSEGLASELSRGMNRMAFDDQSQ
GHRGAQSQEQQGAQSHGQYGALLQRQQEGEAQALSEITAYLQLDDIQTQPITTLASHDVET
CSSNVSQSYSIQELANSSVAGQNSEMSAAIALSLMPNALPPMAGGSTLPVSGASQMLLSNI
QGASLGSNAIDLDQFAASQSQTTQQPSATSQSTIIQSTSILGNHIDQALAEMGLINAQGQS
IQSLAARMPELARHVEQIANQLGLPPADILLKLIENSNRNN

# B

**Codon Optimized Ch-NF-kB**

**ATG**AACCTGCCTGAGGTGACCAAGAACCCTATCTATGGTAGCAGCATGATCGGCGCCAGCTACC
CTATGAGCATGAGCGTCCCCTATCCTTTTAAGGGCGCTGCTAGACTCGAGATCACCAGACAGCC
TAAAAAAAGAGGCTACCGCTTCCGGTACCTGAGCGAGGGCAAGACACACGGCATGCTGCCCGGC
GAGCCTTCTGATTCTGGCGAGAAAGTGTTCCCCAGCGTGCGGATCGTGAATCACCTGGGCAACG
CCAAGGTGGTTATGTACCTGGTGACCGAGGAGGACCCCCAGTTCATCCACCCCCACACCCTGCT
GATCGACAAGATCCCTGTGGGCGGCTACCACATCTTCGACGTTAATGAGGACTTCGACGTGCAG
CTGAAAAACGTGGCTATCCAGCACGTGAACAAGCAGGATCTGCCTACAAAGATGCTGGATAGAC
ACCTGCAGAGCAAGTACATCAAGGAAGTGTACAGCATGAATCAGTACGGAGCCAACCCTAAGAT
CGACATCGACACCTTTGCCTCCAACCTGGCCGACAAAGCCAAAAAGCAGCCTTTCTTCGACCAC
AGAACCAAGCAGGCACTGACCGGCGAGGAAGAAATCCAGCTGTCCCTGGCCATGAGAAAGAAGC
AAAAGGAGATTAACATGAGCAGCTGTAGACTGTGCTTCATCGCCTTCCTGCAGGACCCTGAAAC
CGGCAAGTTCGACAAGATCCTGCAGCCGGTCCATAGCGATCTGATCATCGACGGCAAGAAGAAG
GAAGGCGCCCCTCTGAAGATTATCAGAGTGTCTCACGTGGCCGGATCTGTGGAAGGAGGAAAAG
AAGTGTGGCTGCTGAGCGACAAGATCGATGCCGAGGATACCGAGGTGTACTTCTGGGAACGGAG
CCAGGACAAGCAGGAGATGTTCTGGGAGGGCTTTGGCGAGTTCAACAAGACAGACGTGTACAAG
CAGGCCCTGATTGTGTTTAAGATCCCCCCCTACTGCAACCAGAACATCGATCAGCCCAGAACAG
TGAATCTCCAACTGAGAAGAAAGAAAGACCGGAACTGCGTGTCCGATGTGCATCACTTCAGCTA
CAAGCCTAAGCACTACGACAGATACGGCCTGCGGGAAAAGCGGAGAAAAAACCTGCCACCAGAG
ATCTACGAGAGCACCCCTCCAATCAAGCGGCAACACATGGACTACGCCGCCGATTCCCCTCCTG
CTTCCACAGCCTCTAAGTACCAGCCATATGATAGCGCCCACATCAACAACCCCACCCTGAGCGA
TATCGTGATGCCTCAGGGCAGAAATGAGTACAGAGCCATCGTCGACGCCAGCACCAGGCACTCT
CCTGCCGTGCCCACAACATTCAACGGCCCTCTGGACAACTACCGGAAGAACCACCACAGCTCTA
AAGAAAGCGACACCGACGAGAAC**TGA**

**Codon Optimized Aa-NF-κB**

**ATG**AATATTGGCAATGACATCGATGTGCCCGAGGAACTGCTGCAGCAGATTCTGACCGGCAACT
ACGATTTTAACATGAAGACCGACGACGAGCCTTACCTGGAGATCCTGCTTCAGCCAAAGCAGAG
GGGCTATAGATTCCGCTATAATTCAGAGGGAATCACCCACGGCGGCATTCCTGCCGAAAGCACA
GAGAAGGGAAACAAGAAGTTCCCTACAGTGCACATCGCTAACTATAAAGGGAAGGCTGCCATTT
GCATTACTCTGGTGACCGCTGAGGACCCACCTCAGGTGCATGCCCACAACCTGGTGGGCAAGGA
CGTGACTAAGGGAATGCTGTATTGCGAGGATGACAAGCAGGAATGGGTGAAGTCTTTCACCAAC
CTGGGCATTCAGCACGTGACAAAGAAGGATCTGGTGAAAGTGCTGCATGAAAAGCTGTGCCAGT
CTTATCAGCTGACCCAGTTCTCTGCAGCAATCAAGACTGAGGACGAGACCCAGAGCTTCGACAT
CAGCGCCCTGATCGGCTCCTTTGGCGGCACCGACGGCGCCGGCACCATGATCGACGAGTCCGTG
GCCATGGCCGTGGCCGAGGAGGAGAACAAGAGACTCAGGAAGGAGGCTGAGGAGATGGCAAAGA
CCATCGACCTGAGCGCTGTGAGGCTGTGCATCGTGGCCTACCTGCCTGACGAGGCCGGGCTGCT
GACACGGGCTCTGCCAGCCGTGTTCTCCAAGCCAATCTTCGACCAGAAGGCCGCCCACTCTGGT
CAGCTGAAAATTTGCAGAATTTCCAAGCCAAGTAGCAGCGTGAACGGAGGAGAAGAGGTGTTCC
TGCTCTGCGATAAGGTGCAGAAAGATGACATCTCCGTGAGGTTTTACCAGGAGGATGGAGCCGG

CAACGCCACCTGGGAGGGCTTCGGCAAGTTTACACAGGCCGACGTGCATAAGCAGAGCGCTATT
GTGTTCACCACTCCCCCATACGTGGATAAGGCCATCCAGCGCCCTGTCGAGGTGTGGCTGCAGC
TCAAAAGGGGCAAAGACAAGGAATGCAGCGAACCTGTGAAGTTTACCTATCGGCCCGAGGAGTA
TGACAGGTATAAGATCGGCGAAAAGAGGAGGAAAGGCCTGCCAGAGAACCTGGAGGAGATGCTG
AATACAGCTAAGCAGTCAAGGCCCGCCACCAGCACTATTAGCTCTGCCCCAGCCGATCTGAGCC
AGTTTAACATGCCAGAGGGCGGACCTCTGGACATCTCCGGCATCCGCTTCTTCAAACAGAGCAA
CCAGGGACCTTTTTCTAGCCGGAAG**TGA**

### Codon Optimized Ch-IκB

**ATG**AGCAACCCCAAGGACAAGCTGCCTAACCAGCCCATCGACTCCGGCATCGGAGATTCCTTCG
GCCCCGATTTCGTGCCCGACGACGAGAAGCGGCTGACAGAGGAACTGGAAAACCTGAACATCCA
GCCAAGCAACCAGATCAACACCAAGACACTGCTGGACCAGGCCTTCACCCCTGATGAGGAGAAC
GATACCTACCTGCACGTGTACATCGCCAAGAACAGCCCTGATGAGGCCATGCAGCTGATCGAAA
TGTGCCCTGACAAAAACCTGCTGAACATCCAAAACTATCTCGGCCAGACACCACTGCATGTGGC
CAGCTACGTGAACTTGGATAAGGTGGCTATGTCTCTGGTGCACCACGACGCCAACCTGGAGATT
CAGGACAGAGATGGAAAAAACGTGTTCCACATCTGCGCCGAGAGAGGCCACATCGAGACACTGG
AAAATGTGATCAAGATGGCCTGCCAGACCAACAAGACCAATAGCGCCTGGAACCTGCTGCACAG
CACAGATTTTGAAGGCCAAGCCCCTTTCTTTCTGGCCGCTCTGAACAAGAAAAAGCAGATGTGC
GTGACCCTGGCTAAACTGAATATCGACGTGAACCAGATCGACATCAAGAATGGCAACACCCCTA
TCCACGAGGCCATCCTGGAACAGAACATTGATTACGACTTCCTGGAGTTCCTGGTTAAGACCTG
TAATCTGAACATCAACGCCCAGAACTACGCCGGCATCACAGCCCTGCACCTTGCAGCTGGAAGA
AACGACTCTAATACCTACAGCAAGCTGCTCGAGCTGGGCGCCAATGACTCTTTTGTCGACATCC
GGGACTTCACCCCTGAGATGTGTGGCCCTAGCGACAGCATCCAGCTGCTGAACGGCGTG**TGA**

### Codon Optimized Ch-BCL3

**ATG**CCCAAACCCTACTCCAATCACAGCAAAAAAGCCGAGGAAAACACCGAGCGGTTTCAGAAGG
CCCTCCAATCTAAAATGGCCCAGTCCGTGGCCACACTGAAGACCACCAAGGTGGAAGGACTGTC
CAACCTGGTCGATGCCGCTAGATTTGTTGAGAAGACCAACTCATCCAAAATCACCAAGAAAGCT
AAGCCTAAGCCCATATATAAGGGCAAGGAGAGAGGCAAGGAAAAGAAGTCTCACAGAGAGTCTA
AGCACCTGATCGTGGAAGATACCGACGCCGACGACGAGACTAGCGACATCGAGTCTGAGGTCGA
GAGCGACTCCCAGCTGGCCCTGCTGAAGAAGAGCGTGTCCGAAATGATGGCCAAGATCCAGCGG
CTGGAGAGCAAGAGAAAGCGGAAGAAGATGCCAAAAAAGGCACACCTGAGCGATTACGAGGATA
GCAGCGATGAGGATGTGAAGCCTAAGAAGCCCCTGAAAAAGGAATCAAGCCCTCCTACAAAGCC
CATCAACGCTAACCAGCAGCTGACCATGCACTCTCAACAGATGATGCAGATGCAGCAGCAGTAC
TGGCAGATGCAGCAGATGATGAACATGTACAGCAACCCTTCTATCAACCCCGCCGGCATGATGC
CTAGCATGCCTTTCATGACCACACAGCACCAGCAACAACCTAGCTTCGTCAACATGAACTCTAA
GAGCGCCAGCAACACCCTGCGGTTTCCTAAGACAAAGAAGCCTAACGCTGAAACCAGATTCAAG
GATAGCAGCAAAATTGAGGACAAGAAGTCCAGAAAGAGCGTGGAAGGAGAGGCCGGCGAGGAAA
CAGACGCAAGCGAGGTGTACGCCCCTACAGCCATCAAGGACCCTATCAACGAGTACCTGGGAAA
AGACCCCTTCGTGCCCACCCCTAGCTTCATCAGCTTCGTGGATGACATCACCAAGGATGAGAAG
ACCGTGCAGAGCACAGTGGTGTCTTCTACAAAGCGCGCCGCCGAGACAAGCAAAGTGCCTGGCA
GCGACATCCCTAGAGCCCTGTTTCAGAGAAATGACTCCATCACCAACACCCAACAGATCTCTGC
CACCAACCAGGGCAACCCCAGCTACGCCGGCATCCTGATGAACCTTTTTCCAACAGGCAAGCCT

AACCCTCTGGAAATCAGAGAGAACGACAAGCGGTCCGAGGAAATCAACACAGTGCTGCGGAGCA
TGATCGGCTACAACCCTGTGGACGCCAAGCCAAAGCAGCCTAAGAGATTCGACCTGTCTGCTGA
TAGGCACACCCACAGCAAGCCCCTGTTCCAAAACCTGCTTCCTATCAGCAAACATAGCCAGTCT
CCTATTCCTAGCGACAACGCCCCTGAGAGCAAGCGGTTCGTGGATAACGCTACATGGCTCAGAG
AATGTAGCAAGAAAATGGACCTGCAGGACCAGGAGGGCGACACCATCCTGCACATCCTGATCGC
CAAGGAAGAAACCCCTAAGGCCATCGAGGTGATCAAGAAGATGCAACACCTGCCCAGCCTGGAC
ATCCTGAACGCCCTGGGCCAGACGCCCCTGCACCTGGCCATGTACACCAAGAACCTGCCCGTTG
TGAAAGAGCTGCTGCAGCACGGTTCTGACGTGTCCCTGGTGGACGCCCACGGAAACAACCCTCT
CCATATCGCCTGCGAGGAGAATAGCATCGAGATGCTGGAAATCGTGTTCAACGAGGGCCTGAGC
CACTACCAGACCGCCACCAACACCAGCGCCATGATCACAACCCTGGCCCCGGAATACTTCCACA
TTATCAACGCCAGAAACAATCAGGGCCTCGCAGCCCTGCACATCGCCGCTAAAGAGGACAACGA
GGTGATCATCAAGTTCCTGGGCGAACGGGGCGCCGAAATGAACAATCAGGAGGGCAGAAGCGGC
AATACCCCACTGCTGATCGCCCTTATGAAAAACAACTGGAAGATGGCTTCTTTCCTGCTAGAGA
GATTCGTGAACGTGAACATCCCCAATTTCAGCAATTTCTACCCTCTGCATTTCGCCGTGCAGGG
CAACCACGTGGAAATGGTGAAGGCTCTGCTGAGCAGAGGAGCTGAGATGGGCAGCAGAGCCAAC
GACTATGACCAGAACGCCAATACCACAGAGATCAAGCAGCTGCTGGCCAAAGAAATGAGACGGA
GACGGAGAAACCGGATCAAGCTGGAACCTAAGAGCCAGATG**TGA**

### Codon Optimized Aa-BCL3

**ATG**GGCGACGGCCGCGAAGGCCAGAGCCTGCTGCAGAGGGACAAAGAGGGAGATACCTTTCTGC
ATATCGCCACTGCCAGGAATGAGCACAGCTACATTGAGAAAATCCTGCGGACCGATGTGGCCAA
AGAGCTGATAAACGCCCAGAATAATCACGGACAGACCGCCCTGCACATCGCTACTATTATGGGG
CAGGGCGACCTGGTGAAACTGCTCCTGCAGTATGGCGCCATTCTGCAGCTGCAGAACAACGATG
GCGACACCTCCATTCACCTGGCCACCAAACTGGGCAAAATTGTGTGCCTGATGATCCTGTGCGA
GAACGTGACAAAGGAAATCCTGAACATCGGCAACGACTCTGGGGAAGTGCCTCTGCACCTGGCT
GCTAAGTCAGAGAATCTGCAGGCCTTTGGCCTGTTCCTGCAGAAAGAGGCCGATGTGAAGTTCC
GCAACGAGAAAACTGGCATGACCGCTCTGCACTTTGCTGTCGAGAAGGAGAACAAGGAGATGGT
TGAAAAGCTGCTGGAGCTGGGCGCCGACGTGAACGCACAGGCCACTAACGGCAACACTCCTTTG
CACAGCCTGCTGGGAGCAGAGCAGCGGAATACAGTCAAGCTGCTGCTGGACAAGGGCGCCGACA
TCAACGCCAAGAATGAGGCCGGGCAGAGTCCCCACGATCTGGCAGATAAGACCATGCAGAGGTT
TATGACCCGGAACAGAAAGGACAGAAAGACAAGGAGGCCAAAGAGACCTAAGGGTGCCACTGCT
TCAGACAACCAGCCCGAGCCCGAGACATCCACAACCCCTAAGCCCGTGTCAGACCCCACACATT
CTAACGAAGCACAGCTGTCCAAGAGACCCTTTATTCCCCTGACCAGGGTGCAGAGTCACCCCGG
CAAGAAGCCCAGGCTGTTCGAGGACGGCGGCGCCAGCAACACCAGTCAGGGGTCTGCTGCCCCA
GCTCAGAGACAGCTGACCAGCTCCGCATCCGTGGATCACTGCGTGCAGCTGGACGCATGGGGAC
GCCGGCTGGATAGCCAGAGCCTGAGCAGGGGCAATTCTGATGAGCCCTGCGACGACATCGAGAG
GCAGGAGTGTCTGAAGCTGCCATTCACCGCCTATAAGTCCATGAGCATGCCCGAGTTCAGCAAT
GGGAAGGTGGACGTTACTAACAGCATCTTCGGGCAGCAGCAGCAGACCCTGACCCGGTTCCAAT
CTGAGGGACTGGCCTCCGAGCTGTCCAGGGGCATGAACAGGATGGCCTTTGACGACCAGTCTCA
GGGCCACCGCGGCGCCCAGAGCCAGGAGCAGCAGGGCGCCCAGAGCCATGGGCAGTACGGAGCT
CTGCTGCAGCGCCAGCAGGAGGGAGAAGCCCAGGCCCTGAGCGAGATCACCGCCTATCTGCAGC
TGGACGATATCCAGACACAGCCTATCACAACCCTGGCCTCCCACGACGTTGAAACATGCTCTTC

TAACGTCAGCCAGTCCTACAGTATCCAGGAGCTGGCCAACTCATCAGTGGCCGGGCAGAATAGC
GAGATGTCCGCCGCCATCGCCCTGAGTCTGATGCCAAACGCCCTGCCACCCATGGCCGGCGGCT
CCACCCTCCCAGTGTCCGGAGCCTCCCAGATGCTGCTGTCAAACATTCAGGGAGCCAGTCTGGG
GAGCAACGCCATCGATCTGGACCAGTTCGCCGCCTCTCAGAGCCAGACAACCCAGCAGCCTAGC
GCAACCTCCCAGTCAACCATCATCCAGAGCACAAGCATCCTGGGCAACCACATTGACCAGGCCC
TCGCCGAGATGGGCCTGATCAATGCTCAGGGCCAGAGCATTCAGAGCCTGGCCGCCAGGATGCC
CGAGCTGGCCCGGCATGTGGAACAGATCGCCAACCAGCTGGGCCTGCCCCCAGCCGACATCCTG
CTGAAACTGATCGAAAACAGCAACCGGAATAAT**<u>TAG</u>**

# C

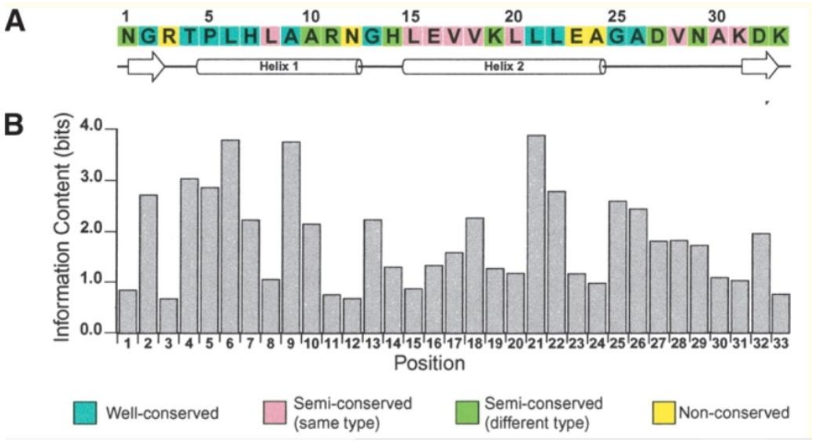


**Supplemental Fig. 1. Amino acid (A) and human cell codon-optimized nucleotide (B) sequences of the proteins used in this study.** In (A), for Ch-IκB, Ch-BCL3, and Aa-BCL3, possible ankyrin repeat sequences are underlined with conserved residues shown in green font and the core TPLH sequences in **bold green font**. In (B), the initiating ATG codons are in **bold font**, and the stop codons for each protein are in **<u>bold underlined font</u>**. In (C), the ANK Repeat Likely Hood Profile used to predict the ANK repeats in the sequences in (B) is shown.

**Supplemental Figure 2**

**Comparison of Aa (top) to Ch (bottom) NF-κB proteins**

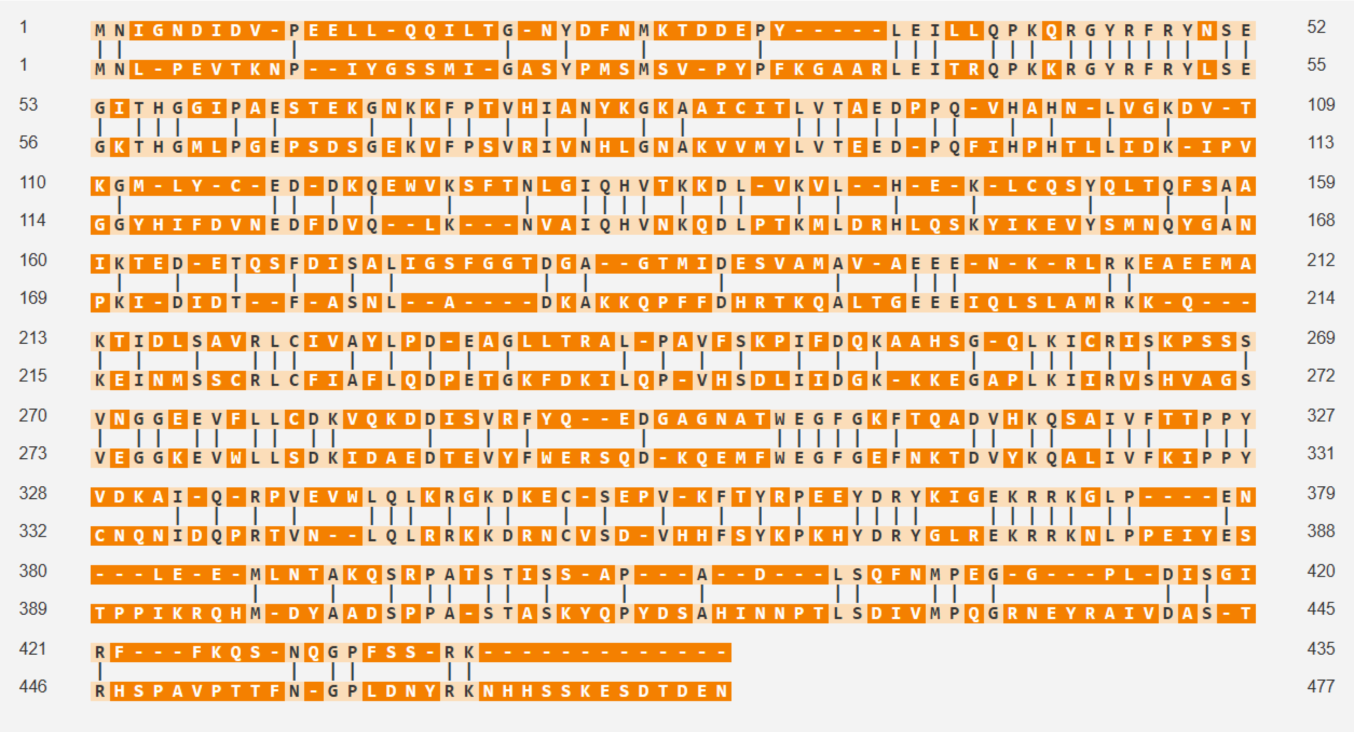


**Supplemental Fig. 2. Comparison of Aa-NF-κB (top) to Ch-NF-κB (bottom).** Comparison was done using the alignment tool at Vector Builder (https://en.vectorbuilder.com/tool/sequence-alignment/252b63da-d27b-4aec-aa9d-d94be64370c1.html). Note the conserved nuclear localization signal (KRRK) at aa 371-374 in Aa-NF-κB and at aa 376-379 in Ch-NF-κB.

**Supplemental Figure 3**

**Ch-IκB** D**S**GIGD**S** **(aa 14-20)**

**Nv-IκB** D**S**GFGG**S** **(aa 41-47)**

**Hs-IκBα** D**S**G-LD**S** **(aa 31-36)**

**Supplemental Fig. 3. Comparison of possible IκB kinase (IKK) serine phosphorylation sites in Ch-IκB to other known IKK phosphorylation sites in IκB proteins.** Shown is an amino acid comparison of the indicated amino acid sequences from human (Hs), *N. vectensis* (Nv) and *C. hemisphaerica* (Ch) IκB proteins. S residues that are conserved phosphorylation sites are in **bold**. S residues in **bold red font** have been experimentally determined to be sites of phosphorylation by IKK protein kinases.

### Supplementary Reference


1. Wolenski, F.S., Garbati, M.R., Lubinski, T.J., Traylor-Knowles, N., Dresselhaus, E., Stefanik, D.J., Goucher, H., Finnerty, J.R., Gilmore, T.D., 2011. Characterization of the core elements of the NF-κB signaling pathway of the sea anemone *Nematostella vectensis*. Mol. Cell. Biol. 31, 1076–1087.
